# Discovery and mechanism of K63-linkage-directed deubiquitinase activity in USP53

**DOI:** 10.1101/2024.07.07.602376

**Authors:** Kim Wendrich, Kai Gallant, Sarah Recknagel, Stavroula Petroulia, Siska Führer, Karel Bezstarosti, Rachel O’Dea, Jeroen Demmers, Malte Gersch

## Abstract

Ubiquitin-specific proteases (USPs) are the largest class of human deubiquitinases (DUBs) and comprise its phylogenetically most distant members USP53 and USP54, which are annotated as catalytically inactive pseudo-enzymes. Conspicuously, mutations in the USP domain of *USP53* cause familial intrahepatic cholestasis. Here we report the discovery that USP53 and USP54 are in fact active DUBs with high specificity for K63-linked polyubiquitin. We demonstrate how USP53 patient mutations abrogate catalytic activity, implicating loss of DUB activity in *USP53*-mediated pathology. Depletion of USP53 increases K63-linked ubiquitination of tricellular junction components. Assays with substrate-bound polyubiquitin reveal that USP54 cleaves within K63-linked chains, whereas USP53 can deubiquitinate a substrate in a K63-linkage-dependent manner. Biochemical and structural analyses uncover underlying K63-specific S2-ubiquitin-binding sites within their catalytic domains. Collectively, our work revises the annotation of USP53 and USP54, provides chemical reagents and a mechanistic framework to broadly investigate K63-polyubiquitin chain length decoding, and establishes K63-linkage-directed deubiquitination as novel DUB activity.

## Introduction

The covalent attachment of the small protein ubiquitin to substrates and to other ubiquitin moieties facilitates the formation of a diverse array of post-translational modifications, which is characterized by structural and functional heterogeneity and collectively termed the “ubiquitin code”.^1,2^ Apart from monoubiquitination, these include polyubiquitin chains of eight linkages with K48 and K63-linked chains being the most abundant in human cells.^1,3^ While attachment of K48-linked ubiquitin chains to substrates commonly facilitates proteasomal degradation, predominantly non-degradative roles have been described for K63-linked chains, including in intracellular trafficking, innate immune signaling and genome maintenance.^4-7^ Ubiquitin chain length encodes an additional layer of information.^1,2,8-14^ However, in contrast to K11- and K48-linked polyubiquitin chains^10,15-19^, molecular mechanisms for length-dependent decoding of K63-linked chains have remained elusive.

Deubiquitinating enzymes (DUBs) edit ubiquitin chains or cleave ubiquitin-substrate linkages, and thereby regulate the ubiquitination status of proteins to determine their stability, localization, activity, and function.^1,20-23^ The human genome encodes approximately 100 DUBs, which can be classified into ubiquitin linkage-specific and non-specific enzymes.^12,16,24-26^ High ubiquitin linkage specificity has been demonstrated for members of the OTU (diverse linkages)^16^, MINDY (K48)^24^, ZUFSP (K63)^7,12,25,27^ and JAMM (K63)^28-30^ families. In contrast, DUBs of the UCH, Josephin and USP families generally show poor discrimination of linkages.^26^ Of those, Ubiquitin-specific proteases (USPs) form the largest and most diverse DUB family with ∼60 members in humans.^31^USP53 and USP54 are poorly characterized USP family members.^32,33^ They possess high homology within their predicted catalytic domains but show otherwise very low sequence homology to all other USP members.^22^ Notably, both proteins have been reported to be catalytically inactive^22,34^ due to the absence of strongly conserved residues before their catalytic histidines^34^, the inability of bacterially expressed USP53 to hydrolyze a mono-ubiquitin substrate^35^ and the lack of reactivity of cellular USP53 with a ubiquitin probe^32^. However, we were intrigued by recent reports which identified loss or biallelic mutations in *USP53* as cause for familial intrahepatic cholestasis, a hereditary liver disorder in children.^36-39^ Notably, this phenotype and proposed cellular roles linked to USP53 and USP54 have not been reconciled with their annotation as pseudo-enzymes.^33^ In addition, the lack of an in vitro assessment of catalytic activity and unavailable structural information have complicated insights into pathology-related molecular mechanisms.

Here we report USP53 and USP54 to be active DUBs with specific cleavage activity on K63-linked ubiquitin chains, revising their annotation as inactive USP family members. Biochemical experiments and a crystal structure of USP54 in complex with a K63-linked diubiquitin probe uncover cryptic S2 ubiquitin sites within the USP domains of both enzymes, underlying efficient cleavage within longer K63-linked chains by USP54. USP53 displays K63-linkage-directed deubiquitination, a DUB activity previously not observed, which is driven by a K63-specific S2 site within its catalytic domain. Taken together, our data establish distinct molecular mechanisms of length-dependent decoding of K63-linked polyubiquitin chains and mechanistically connect loss of USP53 activity to pediatric cholestasis.

## Results

### Activity-based profiling and in vitro assays reveal USP53 and USP54 as active DUBs

Activity-based probes are commonly used for the identification, activity profiling and structural analysis of DUBs.^12,24,25,40^ We recently used propargylamide-based probes^41,42^, whose C-terminal warhead can form a vinyl thioether with catalytic cysteines of active enzymes, to characterize Ub/Ubl cross-reactivity of deubiquitinases (**Supplementary Fig. 1a-b)**.^43^ Consistent with previous studies, the ubiquitin probe facilitated enrichment of many DUBs and HECT family E3 ligases.^40^ Surprisingly, we also detected prominent enrichment of USP54 (**Fig. 1a**) even though this enzyme and its close homologue USP53 are widely reported to be catalytically inactive.^22,32,35^ Based on the catalytic domains, both enzymes are the phylogenetically most distinct members of the USP family of DUBs and cluster with the deSUMOylase USPL1 (**Fig. 1b**).^22^

**Fig. 1.**
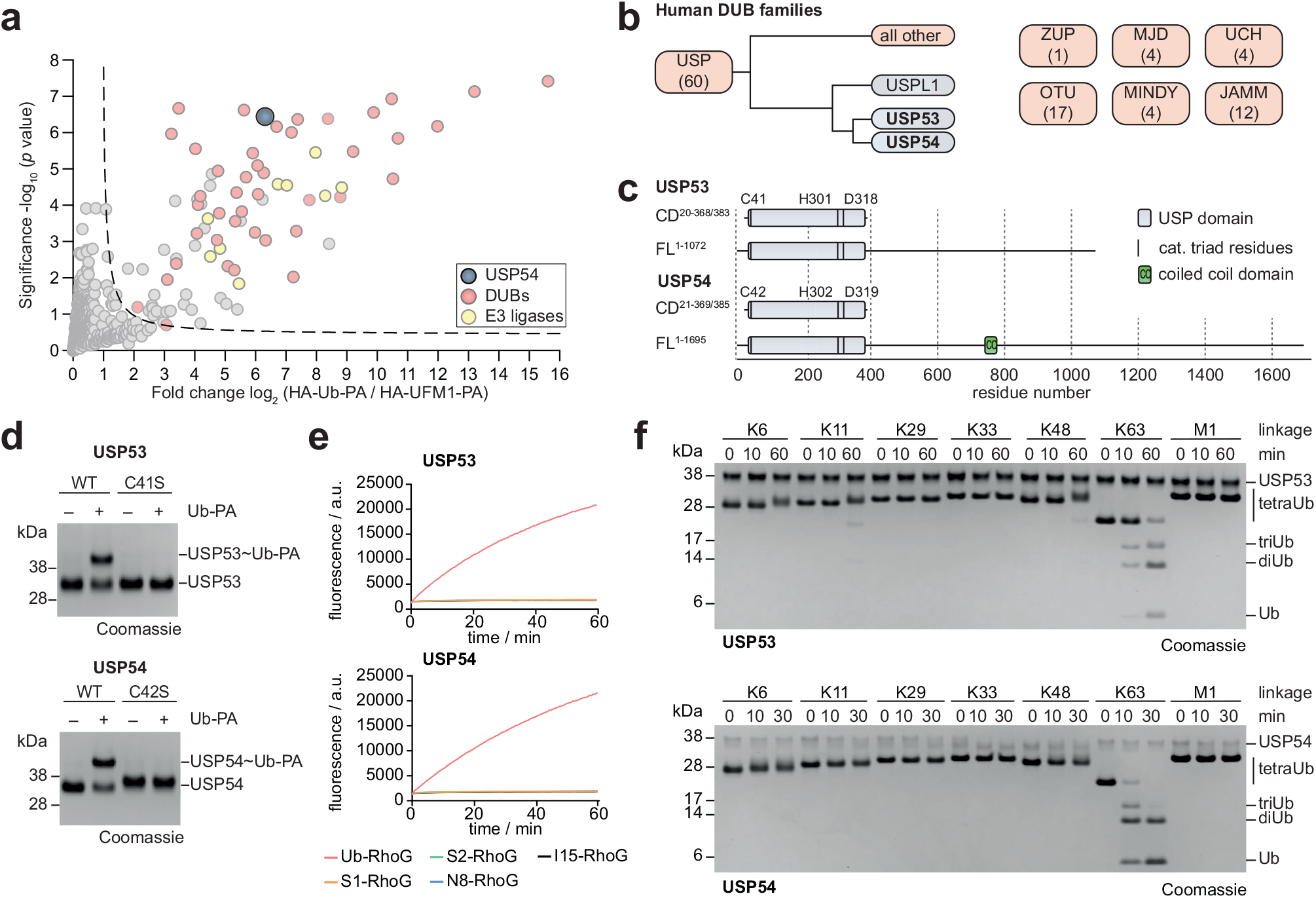
USP53 and USP54 are active DUBs for K63-linked polyubiquitin. **a**. Proteomics analysis of Ubiquitin-PA probe-labelled proteins in HeLa cell lysate. The volcano plot shows the enrichment (fold change) of proteins detected in HA pulldowns from lysate treated with HA-Ub-PA probe in comparison to HA-UFM1-PA as control. Identified proteins are shown as dots, HECT E3 ligases are shown in yellow, DUBs are shown in red and USP54 is shown in blue. **b**. Human DUB families are shown as boxes with number of members given in brackets. USP53 and USP54 were annotated as inactive and comprise the most distantly related catalytic domains within the USP family, with highest similarity to the deSUMOylase USPL1. **c**. Domain architectures of human full-length (FL) and catalytic domain (CD) constructs of USP53 and USP54 used in this study. **d**. Probe reactivity assay with recombinant catalytic domains. HA-Ub-PA was incubated with wild-type USP53^20-383^, USP54^21-369^ and inactive mutants. Probe reactivity was analyzed by SDS-PAGE and Coomassie staining. WT, wild-type. **e**. Ub/Ubl-RhoG cleavage assay. USP53^20-383^ (*top*) and USP54^21-369^ (*bottom*) were added to either Ubiquitin-RhoG (Ub-RhoG), SUMO1-RhoG (S1-RhoG), SUMO2-RhoG (S2-RhoG), NEDD8-RhoG, ISG15-RhoG (I15-RhoG), and fluorescence was recorded. Data are shown as average of technical triplicates. **f**. Gel-based ubiquitin chain cleavage assay. Specifically linked tetraubiquitin chains were incubated with USP53^20-383^ or USP54^21-369^. Samples were taken after indicated time points and cleavage activity was analyzed by SDS-PAGE and Coomassie staining. diUb, diubiquitin. triUb, triubiquitin. tetraUb, tetraubiquitin.

To test for potential activity of USP53 and USP54, we bacterially expressed and purified their catalytic domains (**Fig. 1c**). Using a panel of probes including ubiquitin, different SUMO orthologues and other Ubiquitin-like proteins (Ubls) (**Supplementary Fig. 1c-d**), we observed specific reactivity with HA-Ub-PA for both USP53 and USP54 (**Supplementary Fig. 1c**). This reactivity depended on the presence of the predicted catalytic cysteines (**Fig. 1d**). To directly test for ubiquitin C-terminal hydrolase activity, we employed a panel of Ub/Ubl-derived fluorogenic reagents (**Supplementary Fig. 1e**) and observed concentration-dependent and specific cleavage of Ub-RhoG, but not of SUMO1-RhoG, SUMO2-RhoG, ISG15-RhoG or Nedd8-RhoG, by both USP53 and USP54 (**Fig. 1e, Supplementary Fig. 1f**).

With the notable exception of CYLD, USP DUBs generally show unspecific cleavage or only moderate selectivity towards differently linked ubiquitin chains.^26,44,45^ To analyze cleavage activity on isopeptide linkages we assayed a tetraubiquitin (tetraUb) panel. To our surprise, USP53 and USP54 cleaved K63-linked chains with remarkable specificity (**Fig. 1f**). This finding was highly unexpected as no other human USP DUB shows specific cleavage activity of a single linkage. For USP54, no cleavage was observed for any other linkage whereas USP53 minimally cleaved K11- and K48-linked tetraubiquitin chains at longer time points. Collectively, these experiments demonstrate that human USP53 and USP54 are active enzymes with a specificity for K63-linked polyubiquitin (**Fig. 1, Supplementary Fig. 1**).

### USP53 and USP54 specifically cleave K63-linked, long ubiquitin chains

Recently, 14 reports have linked pediatric cholestasis to biallelic mutations in the *USP53* gene (**Supplementary Table 1)**.^36-39^ However, the falsely presumed enzymatic inactivity of the USP53 protein has so far prevented a mechanistic analysis of the underlying pathologies. We compiled all missense mutations in USP53 and found a clustering within its catalytic domain (**Fig. 2a**). We obtained the USP53 catalytic domain with the disease-associated mutation R99S through bacterial expression and found it to be completely inactive towards K63-linked triubiquitin chains (**Fig. 2b**) and Ub-RhoG (**Supplementary Fig. 1g**). Consistently, USP54 with an equivalent mutation of R100A was virtually inactive in the same assays (**Fig. 2b, Supplementary Fig. 1g**). Both mutant proteins were capable of recognizing ubiquitin, as evident from reduced but retained reactivity towards a ubiquitin probe (**Supplementary Fig. 1h**) and were overall folded as assessed from thermal shift assays (**Supplementary Fig. 1i-j**). These data demonstrate that the disease-associated mutation R99S in USP53 causes a loss of enzymatic activity. More broadly, the data implicate loss of USP53 DUB activity as cause of *USP53*-mediated pathology.

**Fig. 2.**
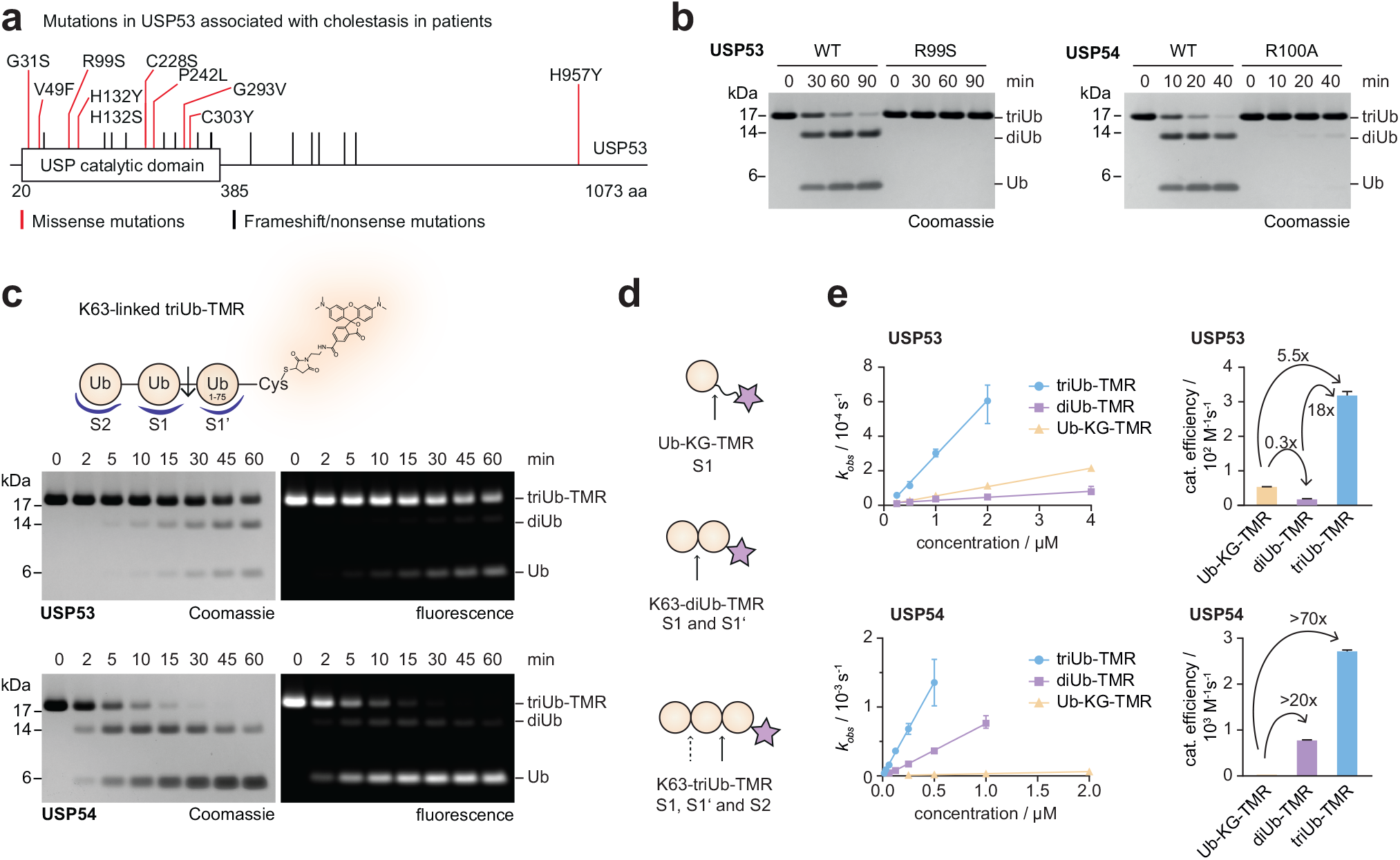
Analysis of USP53 patient mutations and polyubiquitin chain length-dependent cleavage. **a**. Schematic of the human USP53 protein highlighting patient mutations in *USP53* associated with cholestasis or hearing loss. Mutations leading to single amino acid changes are shown in red and cluster in the catalytic domain. Mutations leading to frameshifts or truncations are shown in black. **b**. Gel-based ubiquitin chain cleavage assay. K63-linked triubiquitin chains were incubated with USP53^20-383^ (*left*) or USP54^21-369^ (*right*) with indicated mutations for indicated time points. Cleavage activity was analyzed by SDS-PAGE and Coomassie staining. WT, wild-type. **c**. Fluorescence-based triubiquitin cleavage assay. A K63-linked triubiquitin substrate was used, in which the proximal Ub^1-75^-CA was conjugated to maleimide-TAMRA via the cysteine (triUb-TMR). Cleavage by USP53^20-383^ or USP54^21-369^ was assayed for indicated time points, and analyzed by SDS-PAGE, in-gel fluorescence, and Coomassie staining. The major cleavage position is indicated with an arrow. S2, S1 and S1’ ubiquitin binding sites in the DUB were assigned consistent with the observed cleavage products. **d**. Schematic of ubiquitin substrates used in fluorescence polarization cleavage assays in panel e. Possible cleavage sites are indicated by black arrows. Ubiquitin binding sites (S1, S1’, S2), which when engaged by the DUB lead to cleavage, are given for the respective substrates. TAMRA is shown as a purple star. Structures of the substrates are shown in Supplementary Fig. 2. **e**. Fluorescence polarization cleavage assays. Ub-KG-TMR, diUb-TMR and triUb-TMR were incubated with USP53^20-383^ (*top*) and USP54^21-369^ (*bottom*), and fluorescence anisotropy was recorded over time (see Supplementary Fig. 2 for raw data). Observed rate constants (*k*_*obs*_, shown as mean ± standard error) were plotted over enzyme concentrations to obtain catalytic efficiencies, which are shown for all three substrates in bar graphs as mean ± standard error.

Strikingly, diubiquitin (diUb) chains accumulated during K63-linked polyubiquitin chain cleavage by both enzymes (**Fig. 1f** and **Fig. 2b**), as confirmed in further time-resolved assays (**Supplementary Fig. 2a**). These data suggest an unusual ubiquitin chain length preference of both enzymes. Linkage specificity of DUBs is typically facilitated by S1 and S1’ ubiquitin binding sites, with preferred cleavage of longer chains being facilitated through additional ubiquitin binding sites (**Fig. 2c**).^16,24^ To determine if USP53 and USP54 contain S2’ or S2 sites, we prepared fluorescently labeled, K63-linked triubiquitin (triUb) substrates allowing separate detection of fluorescent species (**Supplementary Fig. 2b-e**). As we found USP54 activity to be impaired by the presence of the FlAsH dye (**Supplementary Fig. 2c**), we generated K63-linked triUb-TAMRA in which the proximal ubiquitin is fluorescently labeled through a C-terminal cysteine (**Supplementary Fig. 2d-e**). Assaying cleavage of the reagent by USP53 and USP54 at early time points revealed preferential conversion into non-fluorescent diubiquitin and fluorescent Ub-TAMRA (**Fig. 2c**). This cleavage pattern suggested the existence of an S2 ubiquitin binding site in both enzymes as depicted in **Fig. 2c** and **Supplementary Fig. 2b**. A potential fourth ubiquitin binding site was excluded through assaying an equivalent K63-linked tetraUb-TAMRA reagent (**Supplementary Fig. 2f-g**).

We next aimed to quantify the contributions of the S1, S1’ and S2 ubiquitin binding sites on USP53 and USP54 activity by using monoubiquitin Ub-KG-TAMRA together with K63-linked diubiquitin and triubiquitin reagents in fluorescence polarization assays (**Fig. 2d, Supplementary Fig. 2h**). Cleavage activity was measured as the decline in anisotropy over time (**Supplementary Fig. 2i**) and was quantified through observed rate constants, yielding catalytic efficiencies of USP53 and USP54 for each substrate (**Fig. 2e**). USP53 showed poor cleavage of Ub-KG-TAMRA and an even lower turnover of the diubiquitin substrate. Addition of a third ubiquitin led to an 18-fold increase in catalytic efficiency (**Fig. 2e**). This behavior could be explained by competing and catalytically unproductive binding of the monoUb and diUb substrates to a high affinity S2 binding site. In contrast, USP54 showed an even poorer cleavage of Ub-KG-TAMRA, but a more than 20-fold increase of catalytic efficiency for diUb-TAMRA. Additional engagement of the S2 site by triUb-TAMRA led to a further increase in catalytic efficiency (**Fig. 2e**), consistent with the results of the tetraUb panel (**Fig. 1f**).

### USP53 displays linkage-directed deubiquitination activity

While DUB activity is typically assayed with free chains, the majority of cellular polyubiquitin is conjugated to protein substrates.^1,2^ To test for distinct activity profiles of USP53 and USP54 in the context of protein-bound ubiquitin chains, we generated a panel of ubiquitylated model substrates including isopeptide-linked monoubiquitylated GFP (GFP-Ub) and GFP conjugated to K48-or K63-linked chains of different length (**Fig. 3a**). The isopeptide linkage of ubiquitin to LACE-tagged GFP was generated with recently reported methodology^46^, and chains were subsequently assembled. Consistent with the fluorescence polarization assays, USP54 did not turnover GFP-Ub, but shortened K63-linked ubiquitin chains to leave a single ubiquitin on GFP (**Fig. 3b**). This finding highlights the importance of its S1’ ubiquitin binding site. Substrate turnover increased with chain length, consistent with the identified S2 site.

**Fig. 3.**
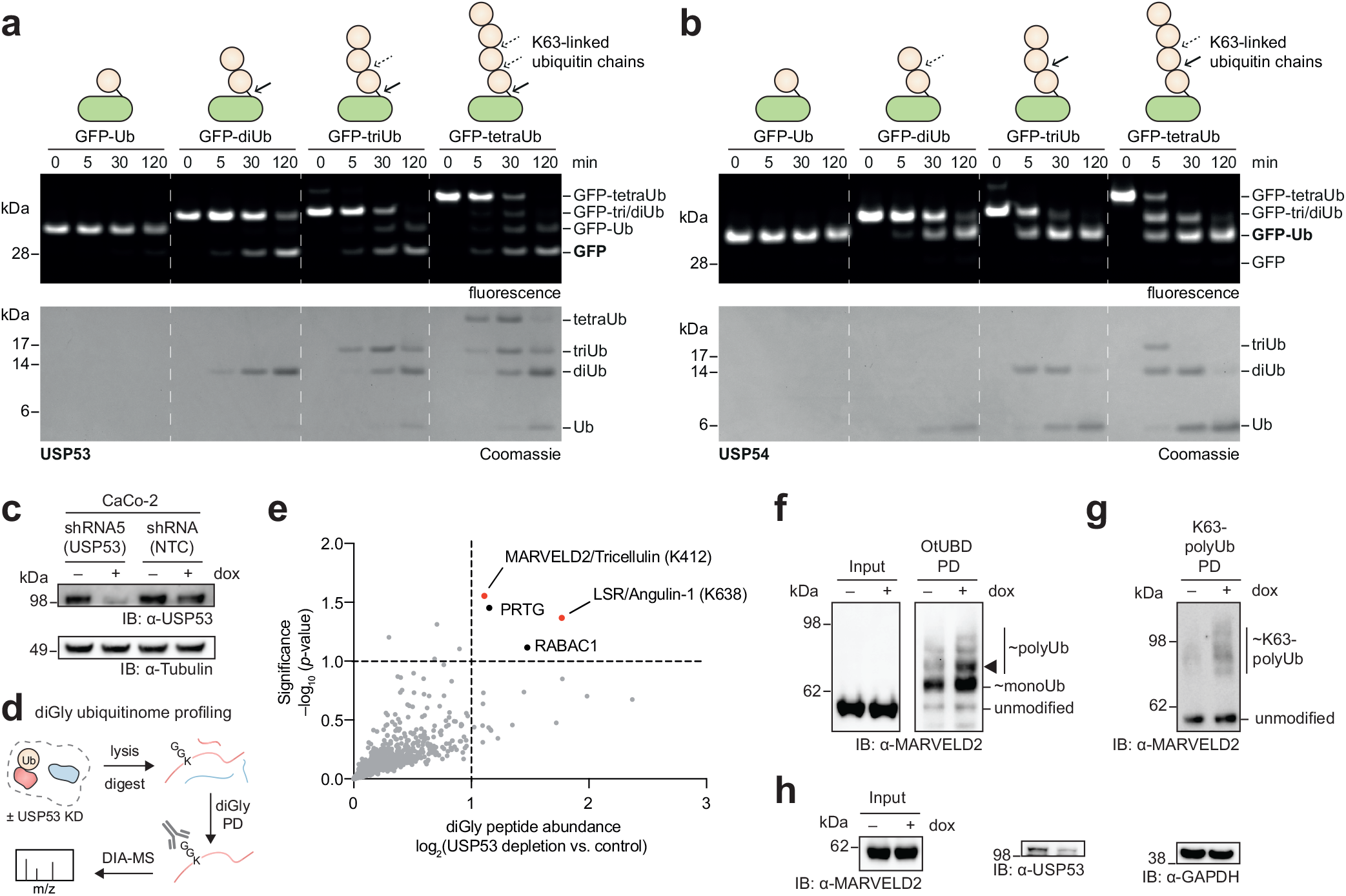
USP53 shows K63-linkage-directed deubiquitination activity. **a-b**. Cleavage assay for isopeptide-linked, mono- or polyubiquitinated model substrates which are modified with K63-linked di-, tri-, or tetraubiquitin chains. GFP-Ub_n_ substrates were incubated for indicated time points with USP53^20-383^ (a) or USP54^21-369^ (b). Cleavage activity was analyzed by SDS-PAGE, in-gel fluorescence, and Coomassie staining. In-gel fluorescence scanning visualized GFP and ubiquitinated GFP proteins, and Coomassie staining of the same gel visualized ubiquitin chains formed during the assay. Observed cleavage events are indicated with arrows. **c**. USP53 depletion analysis. CaCo-2 cells stably transduced as indicated were analyzed by Western blotting after treatment with doxycycline or vehicle control for 72 h. NTC, non-targeting control. **d**. Schematic illustration of the quantitative diGly ubiquitinome profiling experiment to identify potential USP53 substrates. KD, knock-down. PD, pull-down. DIA-MS, data-independent mass spectrometric analysis. **e**. Volcano plot showing the log_2_ fold changes of diGly (ubiquitinated) peptides upon depletion of USP53 in CaCo-2 cells. Significance cut-offs are illustrated with dashed lines. Proteins linked to USP53 phenotypes in patients are indicated in red. diGly modified lysine residues within the peptides are indicated in brackets, with all sites unambiguously identified. **f**. MARVELD2 ubiquitination analysis through OtUBD pull-downs. Cells were treated as in panel G, lysates were enriched with the high-affinity ubiquitin binder OtUBD, and input and PD samples were analyses by Western blotting. The band corresponding to MARVELD2 decorated with two ubiquitin moieties, which emerges upon USP53 depletion, is highlighted with a black triangle. **g**. MARVELD2 polyubiquitination analysis with K63-specific tUIM^Rap80^ (Rx3A7) TUBE pull-down. **h**. Analysis of protein levels for samples processed in panel g.

A strikingly different picture emerged for USP53, which preferentially cleaved polyubiquitin completely off GFP in a K63-linkage dependent manner as evident from gel-based cleavage assays (**Fig. 3a, Supplementary Fig. 3a-d**). We followed the conversion of the GFP substrates by in-gel fluorescence and observed the emergence of free GFP. Coomassie-staining of the same gel visualized free ubiquitin chains generated by USP53. Importantly, GFP-diUb was converted into GFP and free diubiquitin (**Fig. 3a**). The earliest time points of the GFP-triUb and GFP-tetraUb assays showed the emergence of free triubiquitin and tetraubiquitin, respectably, which were later broken down into shorter chains consistent with the tetraubiquitin panel (**Fig. 3a**). The formation of free diubiquitin and triubiquitin chains was validated by intact protein mass spectrometry, which unequivocally confirmed the surprising en bloc deubiquitination activity of USP53 (**Supplementary Fig. 3a-b**). The emergence of low amounts of monoubiquitinated GFP in GFP-triUb and GFP-tetraUb assays demonstrates that substrate deubiquitination is not exclusive and cleavage in substrate-bound chains can take place in K63-linked chains longer than two ubiquitin moieties. Importantly, GFP conjugated to K48-linked diubiquitin or triubiquitin chains was not cleaved under identical conditions (**Supplementary Fig. 3c-d**), demonstrating linkage-specificity of this activity.

Previously reported ubiquitin-linkage-specific DUBs are described to operate strictly within ubiquitin chains. To our knowledge, an enzyme for which ubiquitin chains of a defined linkage direct its deubiquitination activity for the preferred en bloc removal of this modification from substrates has not been described.^2,15,17,18,22,47,48^ Our data strengthen the hypothesis that the en bloc deubiquitination activity of USP53 is facilitated by ubiquitin binding to both the S1 and S2 sites (**Supplementary Fig. 3d**).^22,47^ Taken together, these results show that USP53 and USP54 are active DUBs with intriguing K63 linkage and chain length specificities, with USP53 possessing unique K63-linkage-directed deubiquitination activity.

### Cellular validation of USP53 activity reveals tricellular tight junction components as substrates

To validate this activity in a cellular context and investigate possible consequences related to USP53 loss, we generated stably transduced CaCo-2 cell lines carrying an inducible shRNA against USP53. CaCo-2 is a widely used human model cell line for the study of epithelial cell barriers, and USP53-shRNA induction led to effective depletion of the USP53 protein (**Fig. 3c**). We next assessed the effect of USP53 depletion on the ubiquitination status of proteins by ubiquitinome analysis, carried out by enrichment of diGly-modified peptides and data-independent mass-spectrometric analysis (**Fig. 3d**).^3^ The top four ubiquitination sites, whose abundance increased upon USP53 depletion, included Lys412 of MARVELD2 (also termed Tricellulin) and K638 of LSR (also termed Angulin-1) (**Fig. 3e**). Markedly, these two sites are within the cytoplasmic domains of MARVELD2 and LSR, of which genetic mutations in human patients lead to the same phenotypes as *USP53* mutation.^38,49^ Both membrane-spanning proteins are core components of specialized cell-cell contact sites termed tricellular tight junctions (tTJs),^50^ with LSR recruiting MARVELD2 to tTJs,^51^ and are essential for epithelial barrier function.^49,52,53^ To corroborate ubiquitination of these USP53 substrates, we used the high affinity ubiquitin binding domain OtUBD^54,55^ in a denaturing pulldown of cellular ubiquitin (**Supplementary Fig. 3e-f**). Blotting for MARVELD2 revealed its increased ubiquitination upon USP53 depletion. We observe the strongest change in a species whose molecular weight fits to MARVELD2 modified with two ubiquitin moieties (**Fig. 3f**), which we assign as K63-diubiquitination based on our biochemical data (**Fig. 1f, Fig. 3a** and **Supplementary Fig. 3c**). To directly assess the regulated ubiquitin linkage type, while remaining at endogenous ubiquitin levels, we analyzed the same samples with tandem ubiquitin binding entities (TUBEs) (**Supplementary Fig. 3e-f**).^56^ Their avidity-based mechanism entails that longer chains are enriched much more efficiently than short chains. Pulldown of polyubiquitinated proteins with a high-affinity non-selective 4xUBA^UBQLN1^ reagent showed only a modest increase in overall polyubiquitinated MARVELD2 upon USP53 depletion. However, usage of a K63-specific tUIM^Rap80^ reagent^57^ revealed a strong increase in K63-polyubiquitinated MARVELD2 (**Fig. 3g, Supplementary Fig. 3g-h**). Importantly, global levels of MARVELD2 remained unchanged (**Fig. 3h**), which supports the notion that non-degradative ubiquitination is being regulated by USP53. These results are fully consistent with the biochemically identified K63-linkage-directed deubiquitination activity and validate MARVELD2 as a cellular substrate of USP53.

### A covalent USP54∼K63-diUb-PA complex for crystallization

We next set out to investigate the mechanism of K63 selectivity and how these DUBs recognize chains. To visualize the additional S2 binding site, we aimed to solve a crystal structure of USP53 or USP54 in complex with K63-linked diUb-PA. We prepared this isopeptide-linked probe by enzymatically assembling Ub K63R onto Ub-PA (**Fig. 4a**). A similarly designed K48-linked diubiquitin probe comprising a triazole linkage was previously used for the identification of a K48-specific S2 ubiquitin binding site of a SARS coronavirus PLpro enzyme.^17,58^ We used our probe alongside with Ub-PA to assess ubiquitin engagement in the prepared complexes (**Fig. 4b**). In thermal stability assays, diUb-PA binding stabilized USP53 and USP54 more than Ub-PA, indicating occupation of the S2 site (**Fig. 4c, Supplementary Fig. 4a**).

**Fig. 4.**
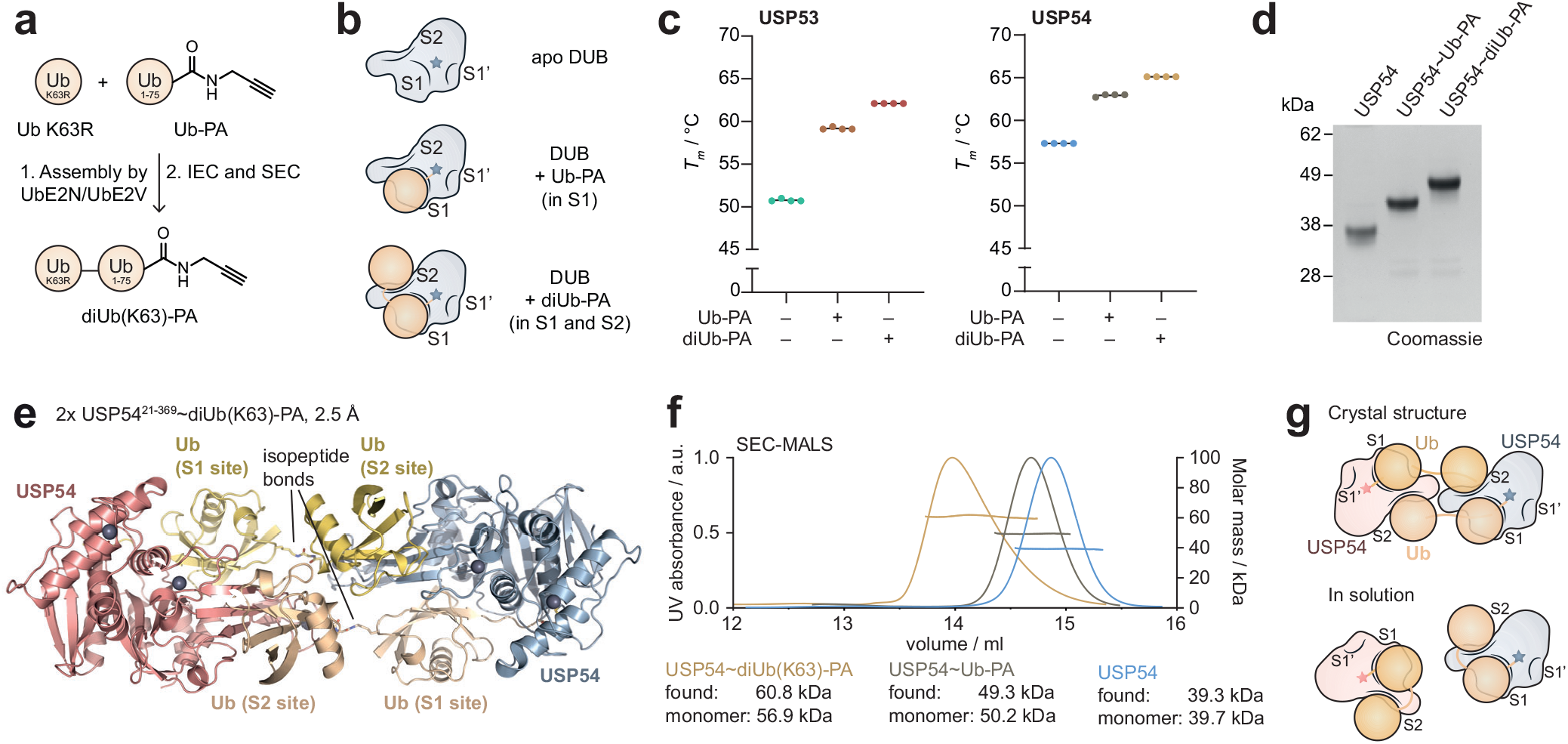
A K63-linked diUb-PA probe enabled crystallization of USP54 in complex with ubiquitin. **a**. Schematic of the generation of a K63-linked diubiquitin probe. The diUb-PA probe was enzymatically assembled from Ub K63R and Ub-PA. IEC, ion exchange chromatography. SEC, size exclusion chromatography. **b**. Schematic of catalytic DUB domains after reaction with Ub-PA or diUb-PA probes, illustrating ubiquitin engagement in samples used in panels c and d. S1’, S1 and S2 Ub binding sites are labelled. The catalytically active cysteine is depicted as a star. **c**. Protein stability assessment. USP53^20-368^ and USP54^21-369^ were labelled with Ub-PA or diUb-PA probes, and stability of protein samples was analyzed by thermal shift analysis. Melting temperatures (*T*_*m*_) are shown for technical replicates, indicating the contribution of the S2 site. **d**. Purified DUB samples. USP54^21-369^, USP54^21-369^∼Ub-PA and USP54^21-369^∼diUb(K63)-PA were analyzed by SDS-PAGE and Coomassie staining. **e**. Crystal structure of USP54∼diUb(K63)-PA. In the observed complex, two USP54 molecules (blue and red) engage two diUb(K63)-PA molecules (gold and wheat) crosswise. The two isopeptide bonds between the diUb-PA molecules are highlighted and are shown as sticks. **f**. SEC-MALS experiment of the catalytic domain of USP54^21-369^ alone (blue), in complex with Ub-PA (brown) or in complex with diUb(K63)-PA (yellow), demonstrating monomeric species in solution. **g**. Schematic depiction of the USP54∼diUb(K63)-PA complex organization in the crystal and in solution.

We prepared and purified several DUB∼probe samples for extensive crystallization trials (**Fig. 4d, Supplementary Fig. 4b-c**). Following construct optimization, we obtained crystals for the most stable complex USP54∼diUb-PA (**Fig. 4c-d**), allowing the determination of a structure to 2.5 Å resolution by X-ray crystallography (**Supplementary Table 2**). Its asymmetric unit contained four copies of USP54∼diUb-PA (**Supplementary Fig. 4d-e**), of which two copies formed an identical arrangement (**Supplementary Fig. 4f**). Excellent electron density covering the entire complex revealed that the two ubiquitin moieties of the probe engaged two different USP54 copies with symmetrical binding *in trans* (**Fig. 4e, Supplementary Fig. 4e**). To assess the solution state of this complex, multi-angle light scattering (SEC-MALS) measurements were performed, proving that all USP54 samples are monomeric in solution (**Fig. 4f**). These results allow the assignment of the dimeric USP54∼diUb-PA arrangement as crystal artefact, likely derived from stabilization of the finger domain of USP54. We propose that in solution the USP54∼diUb-PA complex exists as monomer, and that USP54 engages K63-linked ubiquitin chains through its S1 and the S2 sites *in cis* (**Fig. 4g**).

### USP54 contains a USP catalytic domain with a weakened S1 site

USP54 adopts the USP DUB catalytic domain fold consisting of hand, thumb, and fingers subdomains (**Supplementary Fig. 5a**). A model for the solution state illustrated binding of the distal ubiquitin to an S2 site located at the back of the USP fingers subdomain (**Fig. 5a**). An identical orientation was observed for all four USP54 molecules within the asymmetric unit (**Supplementary Fig. 5b**). Moreover, we found the C-terminal residues of the ubiquitin in the S2 site to be in close proximity to the K63 side chain of the S1 ubiquitin (**Fig. 5a**). This is consistent with an in-solution structure in which the isopeptide bond is placed on top of the fingers without the need of reorientation of the ubiquitin moieties.

**Fig. 5.**
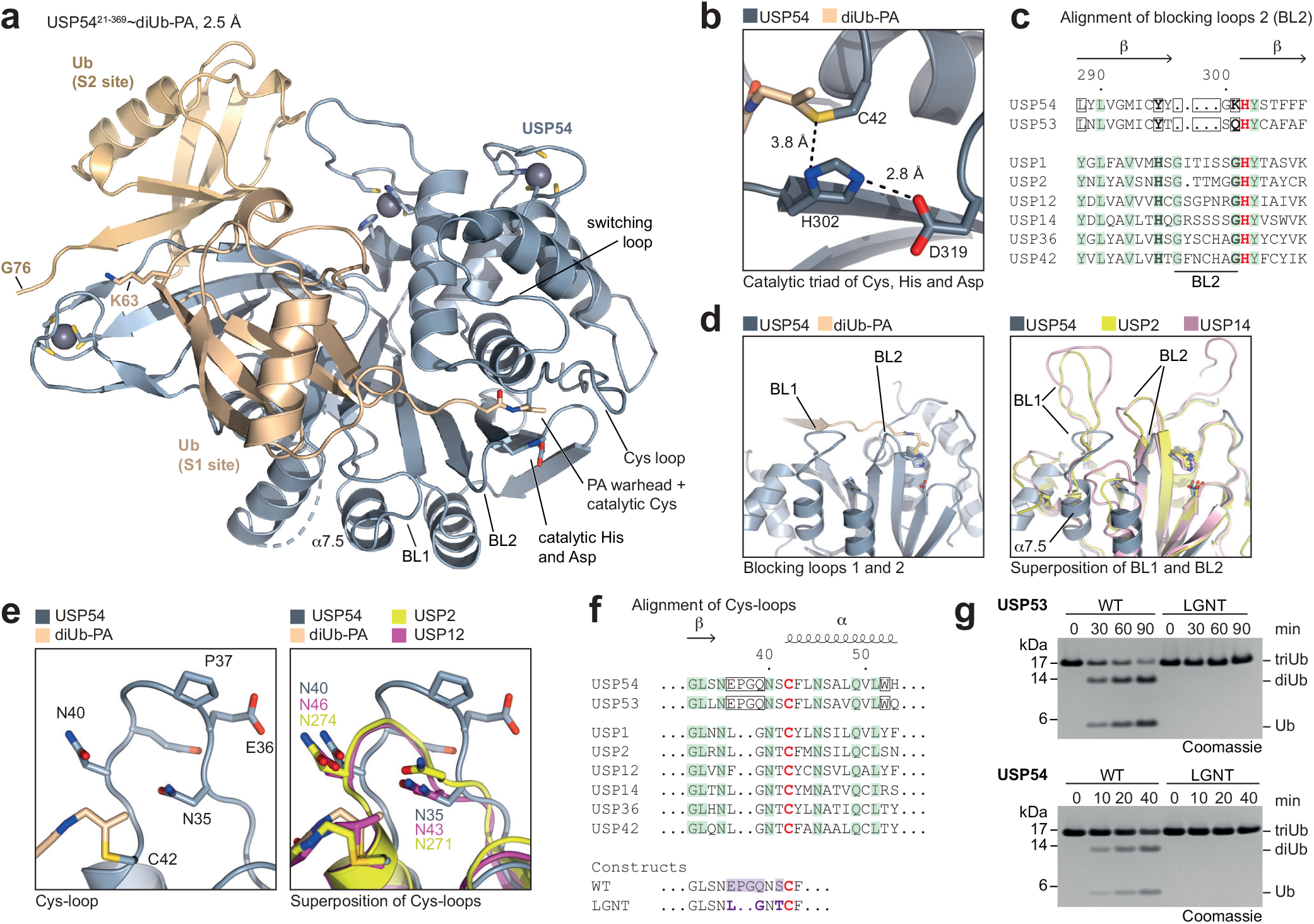
Catalytic activity of USP53 and USP54 depends on a unique Cys-loop. **a**. Solution arrangement of USP54∼diUb(K63)-PA crystal structure. Cartoon representation depicting USP54^21-369^ (grey) with ubiquitin moieties bound in its S1 (wheat) and S2 (gold) ubiquitin binding sites. The K63 side chain and the C-terminal residues of the S1 ubiquitin are displayed as sticks. Residues of USP54 coordinating zinc atoms are shown as sticks, and zinc atoms are shown as grey spheres. The catalytic residues and the residues belonging to the Cys-loop of USP54 are also shown as sticks. BL1, blocking loop 1; BL2, blocking loop 2. 7.5, helix unique in USP54. **b**. Close-up view of the catalytic triad formed by residues C42, H302 and D319. Indicated distances are visualized by dotted lines. **c**. Sequence alignment of residues forming the BL2 in USP DUBs. The catalytic histidines are colored red and conserved residues are colored green. Residues or gaps in USP54 and USP53, which are unique in the entire human USP family, are highlighted with a box. Residues strongly conserved in the USP DUB family, which led to the mischaracterization of USP53 and USP54 as inactive, are additionally highlighted in bold. Secondary structure elements and numbering according to USP54. **d**. Close-up view on BL1 and BL2 of USP54 (*left*). Superposition with the corresponding resides in USP2 (yellow) and USP14 (PDB: 2AYO, cyan), illustrating the atypical shortening of both loops in USP54 (*right*). **e**. Close-up view of the unique loop of USP54 close to the catalytic cysteine (Cys-loop) (*left*). Superposition of the Cys-loop with the corresponding residues in USP2 (yellow) and USP12 (PDB: 5L8W, magenta) (*right*). **f**. Sequence alignment of residues forming the Cys-loop in USP DUBs. The same annotation as in panel c was used. Sequences of USP53 and USP54 constructs used to study the importance of the Cys-loop are shown below with changes marked in violet. **g**. Gel-based ubiquitin chain cleavage assay. K63-linked triubiquitin chains were incubated with USP53^20-383^ and USP54^21-369^ as either WT or Cys loop mutant protein (labelled as LGNT) (see panel f) for indicated time points. Cleavage activity was analyzed by SDS-PAGE and Coomassie staining. diUb, diubiquitin. triUb, triubiquitin.

We next analyzed the USP54 catalytic domain and its engagement of the S1 ubiquitin. A canonical catalytic triad was positioned in an active conformation, with the warhead of the diUb-PA probe covalently linked to the catalytic cysteine (**Fig. 5b**). Cleavage activity on K63-linked ubiquitin chains was abolished for catalytic cysteine and histidine mutants (**Supplementary Fig. 5c-d**). Moreover, the structure revealed three coordinated zinc ions, which in addition to the common position at the tip of the fingers comprise two positions within the thumb subdomain (**Supplementary Fig. 5a**). One of these was identified in human USP36,^43^ whereas coordinating residues for the second zinc are not present in other human USP DUBs.^34^ The positions of the three ions loosely resemble those in the USP domain within the yeast SAGA DUB module.^59,60^

Engagement of the S1 ubiquitin by USP DUBs typically relies on (i.) hydrophobic contacts to the Phe4 patch by the frontside of the finger subdomain, (ii.) a hydrophobic contact of the Ile36 patch by blocking loop 1 (BL1), and (iii.) guiding of the ubiquitin C-terminal residues through the active site cleft by blocking loop 2 (BL2).^43,61^ In USP54, the S1 ubiquitin displayed a ∼30° rotation in comparison to the common binding mode (**Supplementary Fig. 6a**) and differed in all three main interaction sites: i.) The hydrophobic residues normally contacting Phe4 are missing, leaving this patch largely solvent exposed (**Supplementary Fig. 6b**). ii.) Ile36 of ubiquitin is solvent exposed (**Supplementary Fig. 6c**), as BL1 appears absent with corresponding residues adopting a unique helix (α7.5) oriented away from ubiquitin. This arrangement has previously not been observed for USP DUBs (**Fig. 5c-d**). iii.) BL2 of USP54 is truncated (**Fig. 5c-d)** which leaves amide bonds in the ubiquitin C-terminal residues without the otherwise occurring zip-like hydrogen-bonding (**Supplementary Fig. 6d**). Collectively, these differences, which according to the primary sequence appear to be shared by USP53 and USP54, suggest a weakened S1 ubiquitin binding site. This is consistent with the comparably weak activity of both DUBs on monoubiquitin substrates (**Supplementary Fig. 1f, Fig. 2e**).

Another feature of USP53 and USP54, which is unique among human USP DUBs, is an elongated Cys loop directly before the catalytic cysteine (**Fig. 5e-f**). The additional residues are flanked by two asparagine residues conserved in the USP family,^34^ of which Asn35 acts as oxy-anion hole. Mutation of this residue in both USP53 and USP54 verified its importance for catalytic activity (**Supplementary Fig. 5e**). Moreover, mutation of the unique Cys loop sequence (EPGQNSC) of both DUBs into the commonly observed sequence (LGNTC) completely abolished cleavage of K63-linked chains (**Fig. 5g**).

Mapping all cholestasis-related single amino acid mutations in USP53 (**Fig. 2a**) on a homology model illustrated that several mutated residues are either part of the above-described zinc fingers or are located around the active site including R99S (**Supplementary Fig. 6e)**. A closer inspection of Arg99 in USP53 or of the equivalent residue Arg100 in the USP54 structure revealed critical roles in coordinating the unique Cys loop to the switching loop (**Supplementary Fig. 6f-g**), explaining the catalytic inactivity of the mutant proteins (**Fig. 2b**). Taken together, these data demonstrate the importance of the unique active site arrangement in both USP DUBs for catalytic activity and enable a structural rationalization of *USP53*-associated disease mutations (**Supplementary Table 1**).

### A cryptic S2 site within the USP fingers engages K63-linked chains

The structure of USP54∼diUb-PA uncovered an S2 ubiquitin binding site within its USP domain (**Fig. 5a, Fig. 6a, Supplementary Fig. 7a**), consistent with biochemical assay data. This interface is centered on the backside of the fingers subdomain and is governed by hydrophobic interactions with the Ile44 patch of ubiquitin. This main protein interaction site of ubiquitin comprises Ile44 as well as Leu8, Val70 and His68.^1^ The central residue in USP54 at the contact site is Phe161 (**Fig. 6b**), which corresponds to Tyr160 in USP53, suggesting similar ubiquitin engagement. Notably, these hydrophobic residues on the backside of the USP fingers domain are strictly conserved in USP53 and USP54 orthologues (**Supplementary Fig. 7b-c**), whereas other human USP DUBs have varying and typically polar residues at these positions (**Fig. 6c**).

**Fig. 6.**
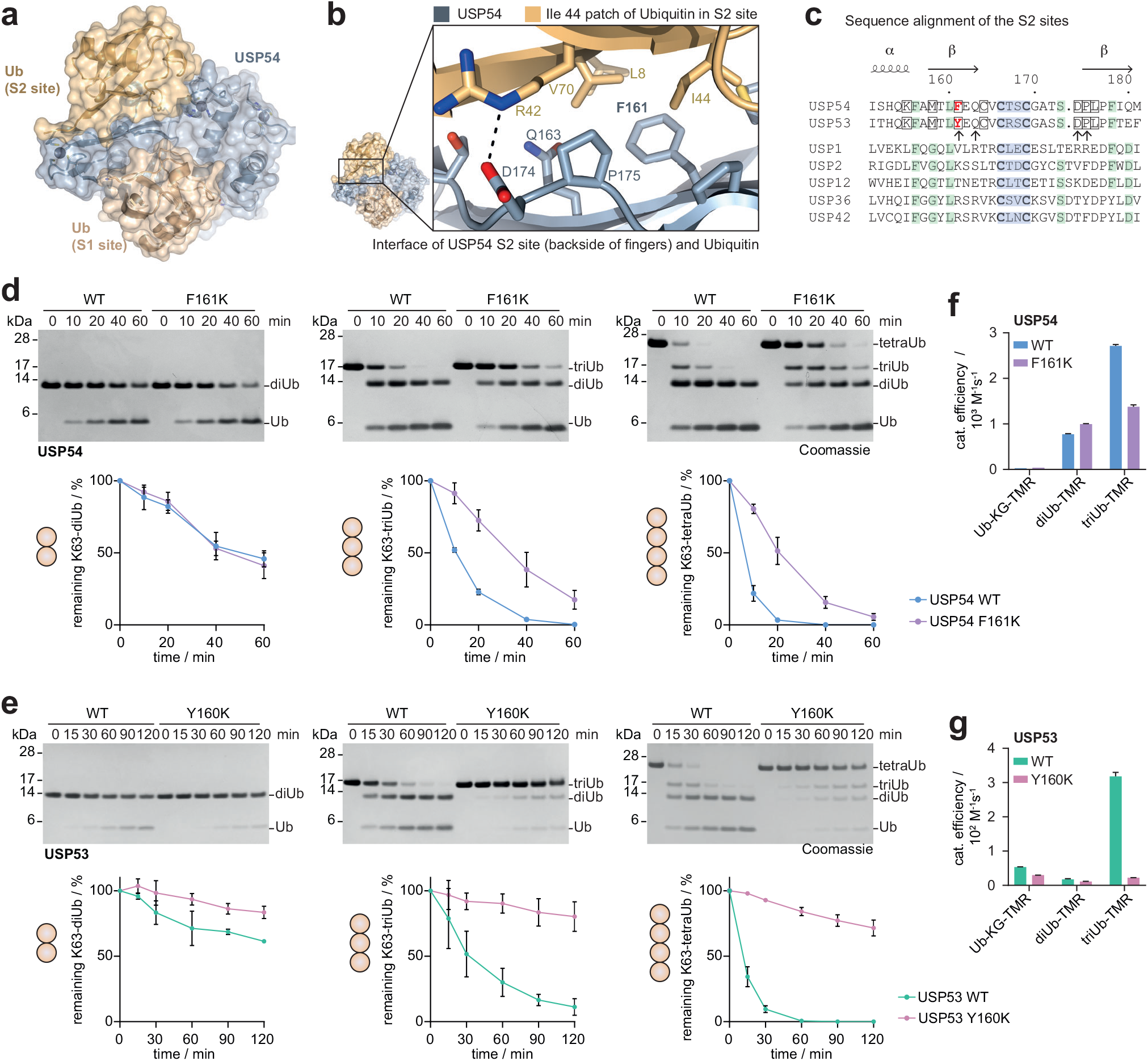
A cryptic S2 ubiquitin binding site in USP53 and USP54 mediates efficient cleavage of K63-linked ubiquitin chains. **a**. Cartoon representation of USP54 catalytic domain in complex with K63-linked diubiquitin. Protein chains are also shown as semitransparent surfaces. **b**. Close-up view on the interface between ubiquitin (gold) and the S2 site of USP54 (grey) located on the backside of the USP fingers subdomain. A hydrogen bond is indicated by a dotted line. **c**. Sequence alignment of residues forming the S2 site in USP54 and USP53, and other USP DUBs. The CxxC motif at the tip of the fingers is marked in blue and conserved residues are colored green. Residues in USP53 and USP54, which are unique in the entire human USP family, are highlighted with a box. Residues annotated in panel B as the S2 site are marked with an arrow. Secondary structure elements and numbering according to USP54. **d**. Gel-based polyubiquitin cleavage assays of USP54. K63-linked ubiquitin chains of different length (di/tri/tetra) were incubated with wild-type USP54^21-369^ or the respective S2 site mutant F161K for indicated times. Cleavage activity was analyzed by SDS-PAGE and Coomassie staining. Substrate consumption was quantified by densitometry of substrate band intensities, normalized to initial intensities. Quantification data are shown as the average ± standard deviation of three independent replicates. **e**. Gel-based polyubiquitin cleavage assays of USP53, as shown in panel d. Wild-type USP53^20-383^ or the respective S2 site mutant Y160K were used. **f**. Catalytic efficiencies of USP54 WT and F161K obtained from fluorescence polarization assays of mono, di, and triubiquitin substrates shown in Fig. 2d. Raw data are shown in Supplementary Fig. 8, and catalytic efficiencies of the WT protein are repeated from Fig. 2e for easy comparison. Data are shown as mean ± standard error. **g**. Catalytic efficiencies of USP53 WT and the S2 site mutation Y160K, analyzed as in panel f. Efficient catalysis of USP53 is dependent on its S2 site, in line with the data shown in panel e.

We next sought to assess the importance of the S2 site for cleavage activity on K63-linked ubiquitin chains. An F161K mutation abrogated binding of the distal ubiquitin through disruption of the S2 site (**Supplementary Fig. 8a-c**). This mutation led to a decrease in cleavage of tri- and tetraUb chains by USP54 (**Fig. 6d**), visualized through quantitative gel-based cleavage assays. Importantly, the mutation did not influence cleavage of K63-linked diUb, demonstrating that binding of Ub chains to the S1 and S1’ sites is not affected by a change in the S2 site (**Fig. 6d**). Strikingly, an equivalent Y160K mutation in USP53 drastically reduced cleavage of tri- and tetraUb chains (**Fig. 6e**), indicating that efficient catalysis for USP53 is driven through the S2 site.

To confirm these findings, we assessed both mutant proteins also in fluorescence polarization assays (**Fig. 6f-g, Supplementary Fig. 8d-e**). Consistently, mutation of either USP53 or USP54 in their S2 sites did not change catalytic activities against monoubiquitin and diubiquitin substrates. However, triubiquitin cleavage activities were dampened. USP53 Y160K showed cleavage activity barely above background (**Fig. 6g**), proving the critical importance of the S2 site for catalysis.

Collectively, these data demonstrate that both USP53 and USP54 possess an additional, K63-linkage specific S2 ubiquitin binding site embedded into their USP catalytic domains. Engagement of the ubiquitin Ile44 patch by the back of their fingers subdomains is unique within the USP family of DUBs, and mechanistically explains their preference for K63-linked ubiquitin chains.

## Discussion

We here revise the annotation of USP53 and USP54 from inactive pseudo-DUBs^34^ to active DUBs, and report in USP53 the discovery of K63-ubiquitin-linkage directed deubiquitination activity. This unprecedented mode of human DUB activity expands the previously established categories of canonical USP DUBs, which typically cleave isopeptide linkages promiscuously, and of linkage-specific DUBs, which edit chains but do not deubiquitinate substrates (**Fig. 7**).^22^ In USP53, recognition of K63-linked chains through an S2 site embedded within its catalytic domain is a prerequisite for efficient catalysis. USP53 shares with its homologue USP54 a weakened S1 ubiquitin binding site through truncations in blocking loops 1 and 2, which explain their comparably poor reactivity on monoubiquitin substrates.

**Fig. 7.**
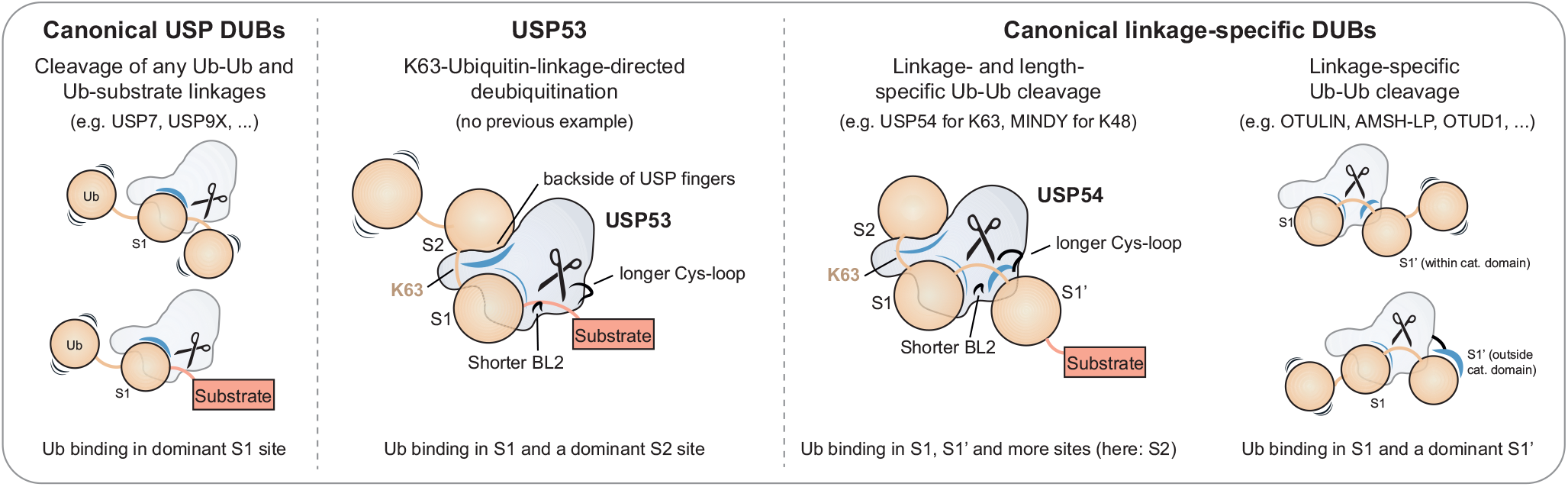
K63-linkage-directed deubiquitination by USP53 bridges canonical DUB categories. Schematic of ubiquitin processing by DUBs. See Supplementary Fig. 9 for a detailed analysis of structural mechanisms for polyubiquitin length and linkage specificity in DUBs of multiple families.

Catalysis driven by an S2 ubiquitin binding site has previously been reported for PLpro enzymes of SARS CoV^17^ and SARS CoV-2^18^, albeit with K48-linkage specificity. These enzymes possess N-terminal extensions to their USP-like catalytic domain to recognize ISG15 and K48-linked polyubiquitin (**Supplementary Fig. 9**), for which hydrophobic recognition of the distal ubiquitin is critical.^62^ While K48-linkage directed substrate deubiquitination by these enzymes has been evaluated *in vitro*,^47,62^ evidence for this activity on cellular substrates is currently not available. Structural^17^ and biochemical^47^ studies on SARS-CoV PLpro concluded that cleavage within K48-linked polyubiquitin is preferred over K48-linkage-directed substrate deubiquitination. This is in contrast to USP53, which when evaluated with purified substrate-bound ubiquitin chains preferred the en bloc removal of K63-polyUb over cleavage within the chain (**Fig. 3a** and **Supplementary Fig. 3a-b**). Moreover, cellular investigation revealed changes in K63-linked ubiquitination of MARVELD2, the emergence of a diubiquitin-modified protein as well as changes in ubiquitination site abundance in MARVDEL2 and LSR upon USP53 deletion (**Fig. 3**). These findings are consistent with the unique activity we characterize, and together set it apart from canonical linkage-specific DUBs. Importantly, deubiquitination by USP53 is greatly accelerated by K63-linked, but not by K48-linked polyubiquitin as expected from the structure-guided biochemical analysis of its K63-specific S2 site (**Fig. 6**). At high concentrations or long incubation times, low cleavage activity on other linkage types can be detected for USP53. The same has also been observed for the M1/K63-specific enzyme CYLD^44^ and is in line with the typically broad catalytic scope of USP family members. This explains why linkage-directed deubiquitination activity could emerge in a USP-fold enzyme with weakened S1 sites and added S2 site. Investigation of a potential S1’ site in future work could illuminate molecular determinants for en bloc substrate deubiquitination versus cleavage within chains.

En bloc removal of ubiquitin chains from substrates has previously been demonstrated for proteasome-resident DUBs PSMD14 and USP14, which is however independent of the chain linkage type.^19,22,48^ Notably, the mechanism of K63-linkage-directed deubiquitination of USP53 is also distinct from the established activity of USP5,^63^ which drives efficient cleavage of free ubiquitin chains through an S1’ site outside of its catalytic and irrespective of the linkage type. Thus, the mechanism of USP53 functionally expands the described DUB activities in human cells.

USP54 is also an atypical USP DUB family member with K63 specificity and a preference for longer chains. Preferred cleavage of longer chains has been reported for: i) ZUFSP for K63-linkages^7,12,25,27^, with a yet unknown mechanism, ii) MINDY DUBs with S2’ through S4’ prime K48-specific Ub sites^15,24^, iii) the Coronavirus PLpro enzyme with a K48-specific S2 site in the N-terminal extension of its catalytic domain^17,18^ and iv) OTUD2 with a K11-specific S2 site.^16^ The mode of ubiquitin chain recognition by USP54 is distinct from all above mentioned examples as well as from the K63-specific DUBs AMSH^28^ and CYLD^44^ (**Supplementary Fig. 8f**), which contain S1’, but not S2 sites within their catalytic domains (**Supplementary Fig. 9**). While K63-linked chains of 2-4 ubiquitin moieties exist on substrate proteins in cells^8^ and can be recognized in a length-dependent manner^13^, how K63 polyubiquitin chain length can be decoded by DUBs on the molecular level was previously not known. These findings will guide the assessment of cellular ubiquitin-dependent roles of USP54 and stimulate research into the chain length-dependent fine-tuning of ubiquitin signaling.

Mutations in *USP53* have clinically been related to bile acids transport disorders, a subgroup of cholestasis^64^ also comprising cases with mutations in tight junction protein 2 (*TJP2*)^65^ and the tricellular junction protein Angulin-1 (*LSR*).^38^ Some patients with congenital *USP53* alterations also presented with hearing loss,^36,39^ consistent with the mouse *USP53* phenotype.^32^ Notably, deafness is also caused by mutations in genes encoding cell-cell junction proteins including claudins, *ILDR1* and the MARVELD family members occludin and tricellulin (*MARVELD2*).^49,53^ USP53 was shown to localize to cell-cell barriers in mouse inner ear cells, and to interact with TJP1 and TJP2.^32^ Consistent with our biochemical and structural data and the proposed model for USP53 activity, we find elevated ubiquitination on the two core components of tricellular junctions MARVELD2 and LSR upon knock-down of USP53 (**Fig. 3c-h**). The combined use of diGly proteomics and different ubiquitin enrichment techniques validates these as USP53 substrates, and synergizes with recent advances in the assessment of ubiquitin chain architecture^3,8,66^ These substrates are supported by coinciding patient phenotypes of deafness and cholestasis,^38,49^ consistent with the *USP53* mouse phenotype.^32,49^ Collectively, the results converge into a model in which loss of the here reported USP53 catalytic activity causes pediatric cholestasis.

Our work critically guides the investigation of ubiquitin-dependent cellular roles of the additional active DUBs USP53 and USP54. Our structural insights will aid the molecular investigation of both enzymes and provide a framework for the understanding of *USP53* disease mutations. Moreover, our results more broadly demonstrate how ubiquitin chains of a particular chain length and linkage can uniquely lead to complete protein deubiquitination by a human DUB in a chain-linkage dependent manner. They stimulate the re-evaluation of known DUBs with isopeptide-linked polyubiquitin substrates – to represent endogenous structures more closely and in addition to frequently used panels of free ubiquitin chains – in search for perhaps other enzymes featuring linkage-directed deubiquitination activity.

## Acknowledgments

We thank the beamline scientists at the Swiss Light Source (SLS) for support during data collection and R. Gasper and P. Geue for support with crystallization and biophysics. We thank all staff at the Max-Planck-Institute for Molecular Physiology for excellent technical support and P. Janning for mass spectrometry support. We are grateful to all members of the Gersch lab for discussions, advice, and reagents. We thank L.-M. Kaps, J.A. Hane and M. Reichmann for support with biochemical assays and cell culture. We are grateful to D. Komander (MRC LMB Cambridge / WEHI Melbourne), D. Alessi (MRC PPU Dundee), D. Trono (EPFL) and D. Wiederschain for the gift of plasmids. This work was funded by AstraZeneca, Merck KGaA, Pfizer Inc., and the Max Planck Society as part of the Chemical Genomics Centre III. Work in the Gersch lab is further supported by Deutsche Forschungsgemeinschaft (DFG, German Research Foundation, Project-ID GE 3110/1-1 – Emmy Noether; and Project-ID 424228829 – SFB1430) and by the State of North Rhine-Westphalia through the ‘CANcer TARgeting Network’ initiative (NW21-062C).

## Author Contributions

K.W. and M.G. designed the study. K.W. performed all experiments unless noted otherwise. K.G. performed cellular experiments. K.W. and S.R. performed *in vitro* assays with mutant proteins. S.P. generated cell lines. S.F. provided Ub enrichment reagents. R.O.D. contributed data in Fig. 1a and Supplementary Fig. 1d. K.B. and J.D. performed ubiquitinome analysis. All authors designed experiments, analyzed data, and interpreted results. M.G. supervised the project and wrote the manuscript together with K.W. and with input from all authors.

## Declaration of Interests

The authors declare no competing interests.

---

**Supplementary Table 1.**
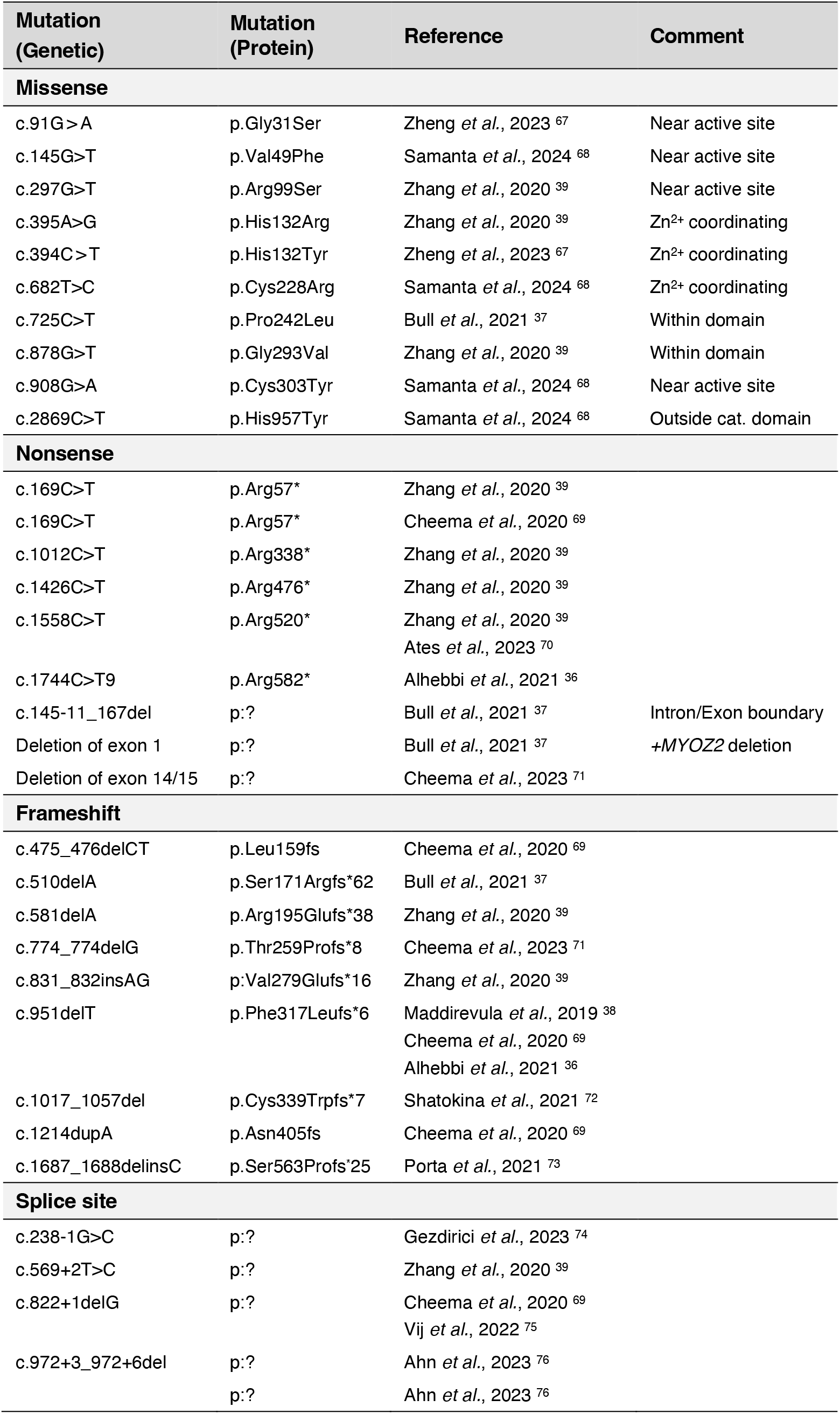
Patient mutations in *USP53* associated with familial intrahepatic cholestasis.

**Supplementary Table 2.** Data collection and refinement statistics.

|  | USP54~diUb(K63)-PA<br>(SAD data) | USP54~diUb(K63)-PA<br>(native data, PDB code: 8C61) |
| --- | --- | --- |
| **Data collection** |  |  |
| Beamline | SLS – PX2 | SLS – PX2 |
| Wavelength | 1.282 Å | 1.000 Å |
| Space group | *P* 2<sub>1</sub>2<sub>1</sub>2<sub>1</sub> | *P* 2<sub>1</sub>2<sub>1</sub>2<sub>1</sub> |
| Cell dimensions |  |  |
| *a*, *b*, *c* (Å) | 122.58, 126.59, 144.22 | 122.74, 126.56, 144.17 |
| $\alpha$, $\beta$, $\gamma$ (°) | 90, 90, 90 | 90, 90, 90 |
| Observed reflections | 1,859,182 (122,127) | 699,375 (71,819) |
| Unique reflections | 69,709 (4,449) | 78,247 (7,759) |
| Resolution (Å) | 88.06 – 2.6<br>(2.66 – 2.6) | 63.28 – 2.5<br>(2.59 – 2.5) |
| *R*<sub>merge</sub> | 0.091 (2.410) | 0.062 (1.239) |
| *R*<sub>meas</sub> | 0.094 (2.498) | 0.066 (1.313) |
| *I*/$\sigma$(*I*) | 24.5 (2.2) | 16.2 (1.7) |
| *CC*<sub>1/2</sub> | 1.000 (0.789) | 0.998 (0.682) |
| Completeness (%) | 100 (100) | 100 (100) |
| Redundancy | 26.7 (27.5) | 6.6 (6.3) |
| Wilson *B* (Å<sup>2</sup>) | 74.4 | 74.8 |
| **Phasing** |  |  |
| Method | SAD | MR |
| Resolution (Å) | 2.6 |  |
| Anom. completeness (%) | 100 (100) |  |
| Anom. multiplicity | 13.8 (13.9) |  |
| <FOM> | 0.5179 |  |
| **Refinement** |  |  |
| Copies / a.s.u. |  | 4 |
| Resolution (Å) |  | 2.5 Å |
| No. reflections |  | 78,226 |
| *R*<sub>work</sub> / *R*<sub>free</sub> (%) |  | 20.0 / 24.3 |
| No. atoms |  | 14,892 |
| Protein |  | 14,647 |
| Ligand |  | 32 |
| Water |  | 213 |
| *B* factors (Å<sup>2</sup>) |  | 95.9 |
| Protein (Å<sup>2</sup>) |  | 96.2 |
| Ligand (Å<sup>2</sup>) |  | 88.8 |
| Water (Å<sup>2</sup>) |  | 79.5 |
| R.m.s.d. |  |  |
| Bond lengths (Å) |  | 0.004 |
| Bond angles (°) |  | 0.62 |
| Ramachandran (favored /<br>allowed / outlier) (%) |  | 96.8 / 3.2 / 0 |
| Clashscore |  | 8.5 |
| Rotamer outliers (%) |  | 2.8 |

The dataset was collected from a single crystal. Values in parentheses are for highest-resolution shell. a.s.u., asymmetric unit. R.m.s.d., root mean square deviations.

**Supplementary Fig. 1.**
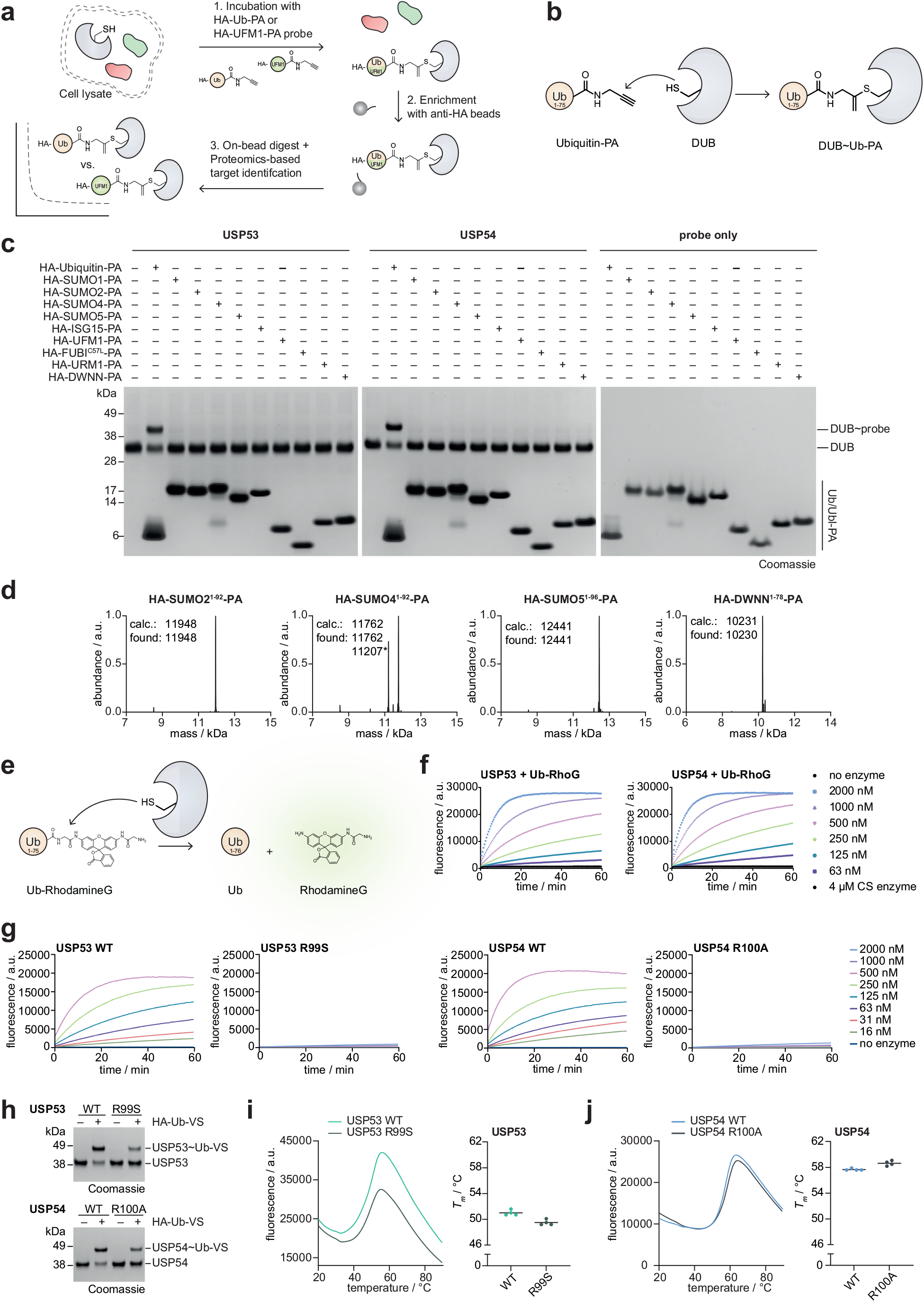
USP53 and USP54 are active DUBs. **a**. Schematic of the workflow to identify active DUBs. HeLa cell lysate was treated with HA-Ub-PA or HA-UFM1-PA probe. Labelled proteins were enriched and detected by proteomics. **b**. Schematic of the probe reactivity assay. The C-terminal propargylamine (PA) warhead reacts with the catalytic cysteine of DUBs to a vinyl thioether, forming a covalent DUB∼Ub-PA complex. **c**. Ub/Ubl specificity assayed through probe reactivity with recombinant catalytic domains. A panel of HA-Ub-PA and HA-Ubl-PA probes was incubated with wild-type USP53^20-383^ and wild-type USP54^21-369^. Probe reactivity was analyzed by SDS-PAGE and Coomassie staining. **d**. Intact protein mass spectrometry of indicated probes used in panel c. Expected and found masses are given in Dalton. The star denotes an HA-SUMO4^1-92^ species N-terminally truncated by four residues. All other probes have been characterized previously.^43^ **e**. Schematic of the Ubiquitin-RhodamineG (Ub-RhoG) cleavage assay. DUBs cleave the amide bond between ubiquitin and RhoG and thereby liberate fluorescent RhoG. **f**. Ub-RhoG cleavage assay. Indicated concentrations of USP53^20-383^ (*left*) and USP54^21-369^ (*right*) or of the catalytic inactive mutants were added to Ub-RhoG and fluorescence was recorded. Data are shown as average of technical triplicates. CS enzymes denotes the use of catalytic cysteine to serine mutated enzymes as in Fig. 1d. Of note, this activity is approximately 3 to 4 orders of magnitude weaker than for other USP DUBs including USP2, USP7, USP36 and USP42.^43^ **g**. Ub-RhoG cleavage assay. Indicated concentrations of WT USP53^20-368^ (*left*) and USP54^21-385^ (*right*) or of the indicated mutants were added to Ub-RhoG and fluorescence was recorded. Data are shown as average of technical triplicates. **h**. Probe reactivity assay. HA-Ub-VS was incubated with proteins shown in Fig. 2b, and probe reactivity was analyzed by SDS-PAGE and Coomassie staining. **i**. Protein stability assessment of USP53^20-368^ WT and USP53^20-383^ R99S by thermal shift analysis. Fluorescence raw data are shown and melting temperatures (*T*_*m*_) are plotted from technical replicates. **j**. Protein stability assessment of USP54^21-369^ WT and USP54^21-369^ R100A by thermal shift analysis as described in panel i.

**Supplementary Fig. 2.**
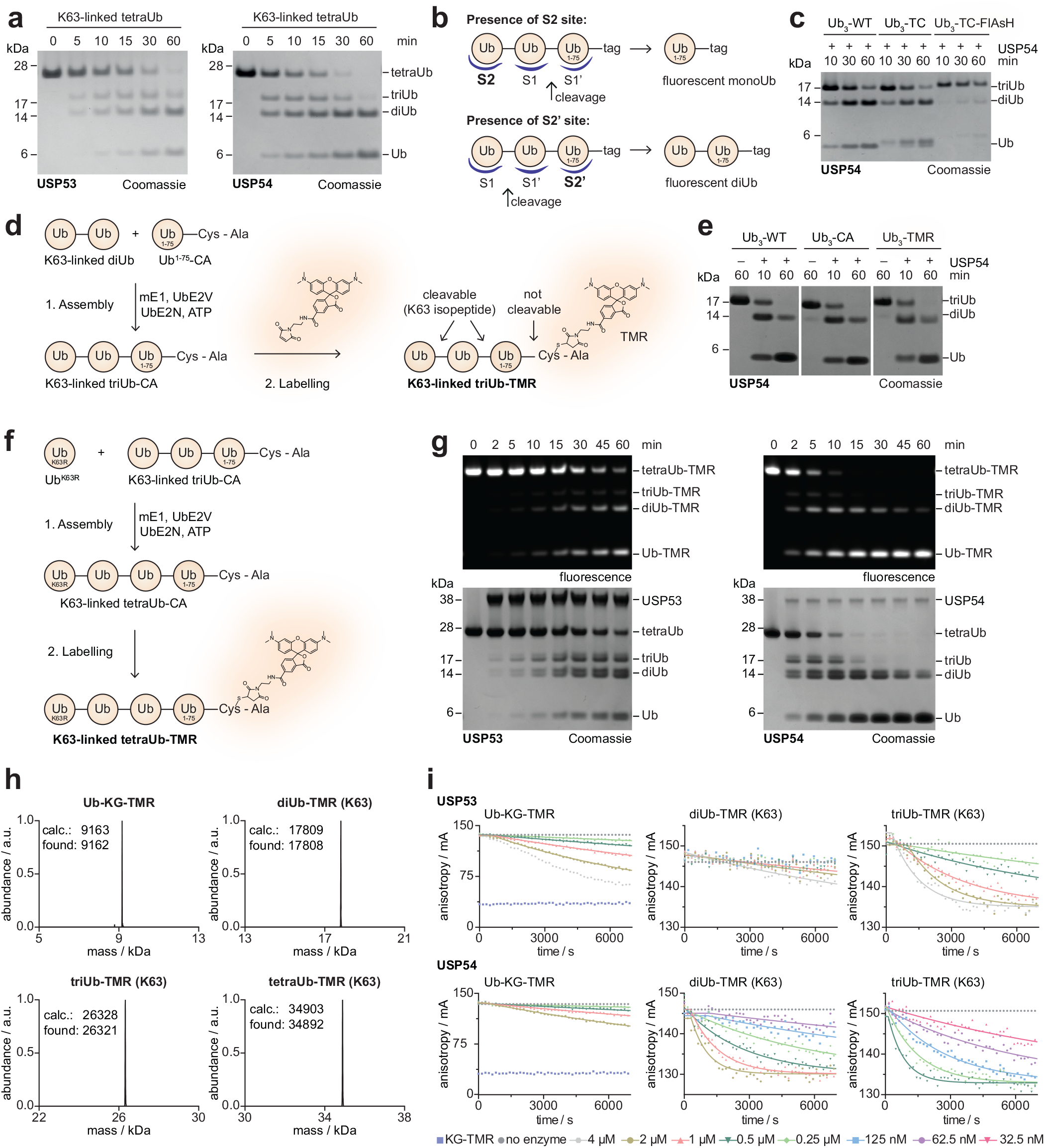
An S2 site, but not an S3 site, underlies USP53 and USP54 polyubiquitin cleavage. **a**. Time-resolved gel-based cleavage assays of K63-linked tetraubiquitin by USP53^20-383^ (*left*) and USP54^21-369^ (*right*). Cleavage activity was analyzed by SDS-PAGE and Coomassie staining. **b**. Schematic of expected products of a triubiquitin cleavage assay in which the reagent contains a non-cleavable, C-terminal fluorescent label. Ubiquitin binding sites in a DUB are shown in blue, the expected cleavage site is indicated with a black arrow. The presence of an S2 site would lead to fluorescently labeled monoubiquitin, whereas the presence of an S2’ site would lead to fluorescently labeled diubiquitin. **c**. Gel-based cleavage assay with a FlAsH-based reagent. Native K63-linked triubiquitin chains, K63-linked triubiquitin with a C-terminal tetracysteine (TC) tag in the proximal Ubiquitin (Ub_3_-TC) and FlAsH-labeled triubiquitin (Ub_3_-TC-FlAsH) were incubated with USP54^21-369^ for indicated times. USP54 activity was impaired by the presence of the FlAsH dye. **d**. Schematic of the generation of the TAMRA-based, fluorescent K63-linked triubiquitin substrate. TMR, TAMRA. **e**. Gel-based cleavage assay with a TAMRA-based reagent. Equivalent assays as shown in panel c with K63-linked triubiquitin chains shown in panel d. **f**. Schematic of the generation of the TAMRA-based, fluorescent K63-linked tetraubiquitin substrate. **g**. Gel-based cleavage assay. Fluorescent K63-linked tetraubiquitin was incubated with USP53^20-383^ and USP54^21-369^ for indicated time points. Cleavage activity was analyzed as in Fig. 2d. The equally formed fluorescent diubiquitin and monoubiquitin species demonstrate that neither USP53 nor USP54 possess an S3 ubiquitin binding site. **h**. Deconvoluted intact protein mass spectra of indicated reagents. Calculated and observed molecular weights are given in Dalton. **i**. Fluorescence polarization cleavage assays of USP53^20-383^ and USP54^21-369^ and indicated reagents. Catalytic efficiencies derived from these data are shown in Fig. 2e.

**Supplementary Fig. 3.**
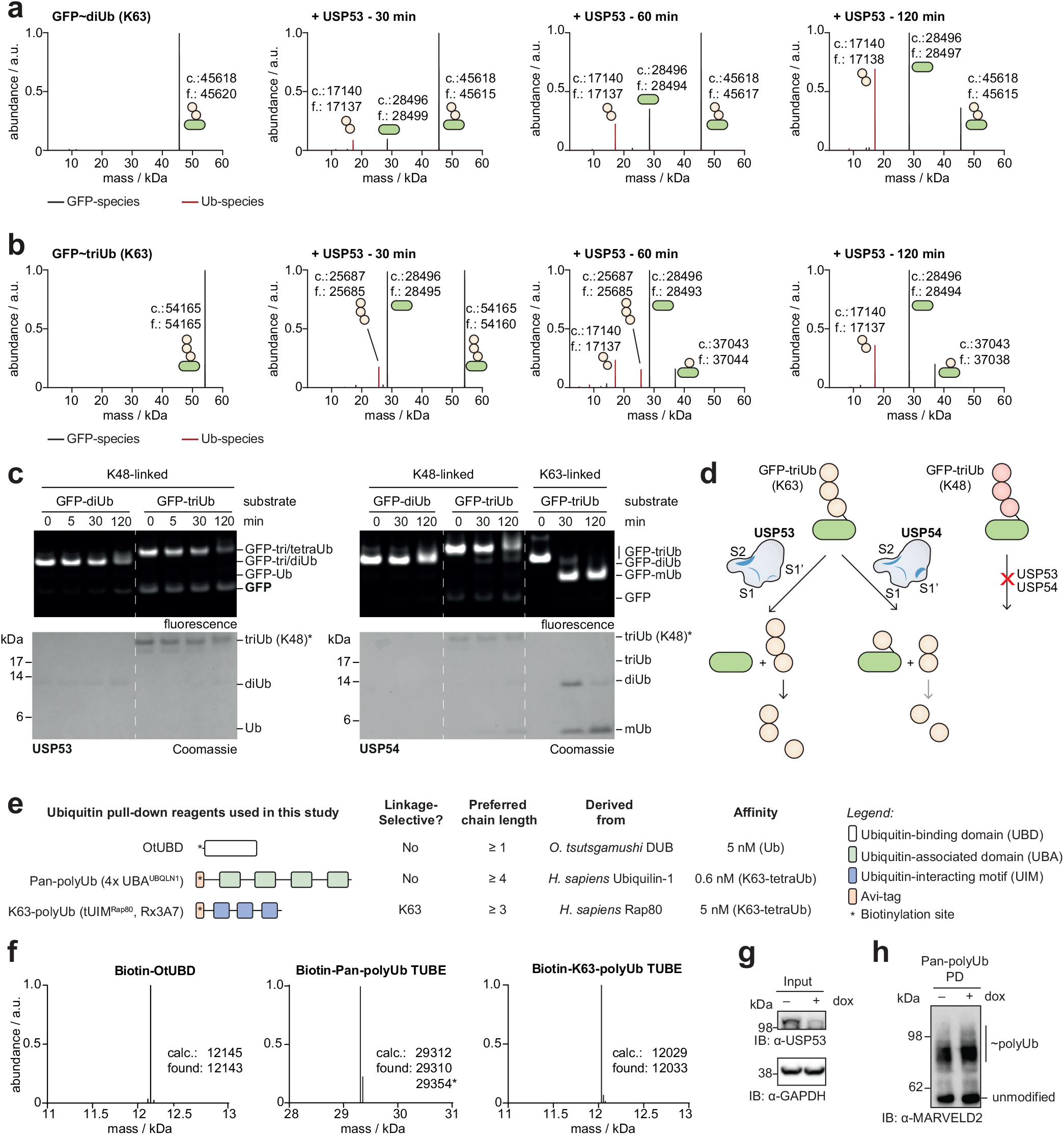
Validation and specificity analysis of linkage-directed deubiquitination activity in USP53. **a**. Intact protein mass spectrometry analysis of GFP-diUb (K63) cleavage assay, corresponding to Fig. 3b. Diubiquitin removed en bloc by USP53 was identified unequivocally by its mass. Calculated (c.) and found (f.) protein masses are given in Dalton. **b**. Intact protein mass spectrometry analysis of GFP-triUb (K63) cleavage by USP53, shown as in panel a. Of note, triubiquitin removed en bloc during the earlier time points was cleaved further to diubiquitin. While en bloc removal was observed exclusively in the earliest time point, cleavage within the chain leading to accumulation of a minor amount of monoubiquitinated substrate was observed at later time points, both consistent with the gel-based assay shown in Fig. 3b. **c**. Cleavage assay for ubiquitinated model substrates modified through isopeptide bonds with K48-linked di- or triubiquitin or with K63-linked triubiquitin. GFP-Ub_n_ substrates were incubated with USP53^20-383^ or USP54^21-369^. Cleavage activity was analyzed by as shown in Fig. 3a-b. **d**. Schematic of the identified cleavage activities of USP53 and USP54 on GFP modified with K63-or K48-linked triubiquitin chains. **e**. Overview of ubiquitin pull-down reagents used in this study. Data are derived from the original publications.^54,56,57^ **f**. Deconvoluted intact protein mass spectra of biotinylated reagents. Calculated and observed molecular weights are given in Dalton. The species in the Pan-polyUb TUBE denoted with an asterisk and a mass difference of 42 Da corresponds to a non-covalent acetonitrile adduct formed during analysis. **g**. Input protein levels for samples used in Fig. 3f and panel h, and analyzed as described for Fig. 3h. **h**. MARVELD2 polyubiquitination analysis with pan-polyubiquitin (4x UBA^UBQLN1^) TUBE pull-downs.

**Supplementary Fig. 4.**
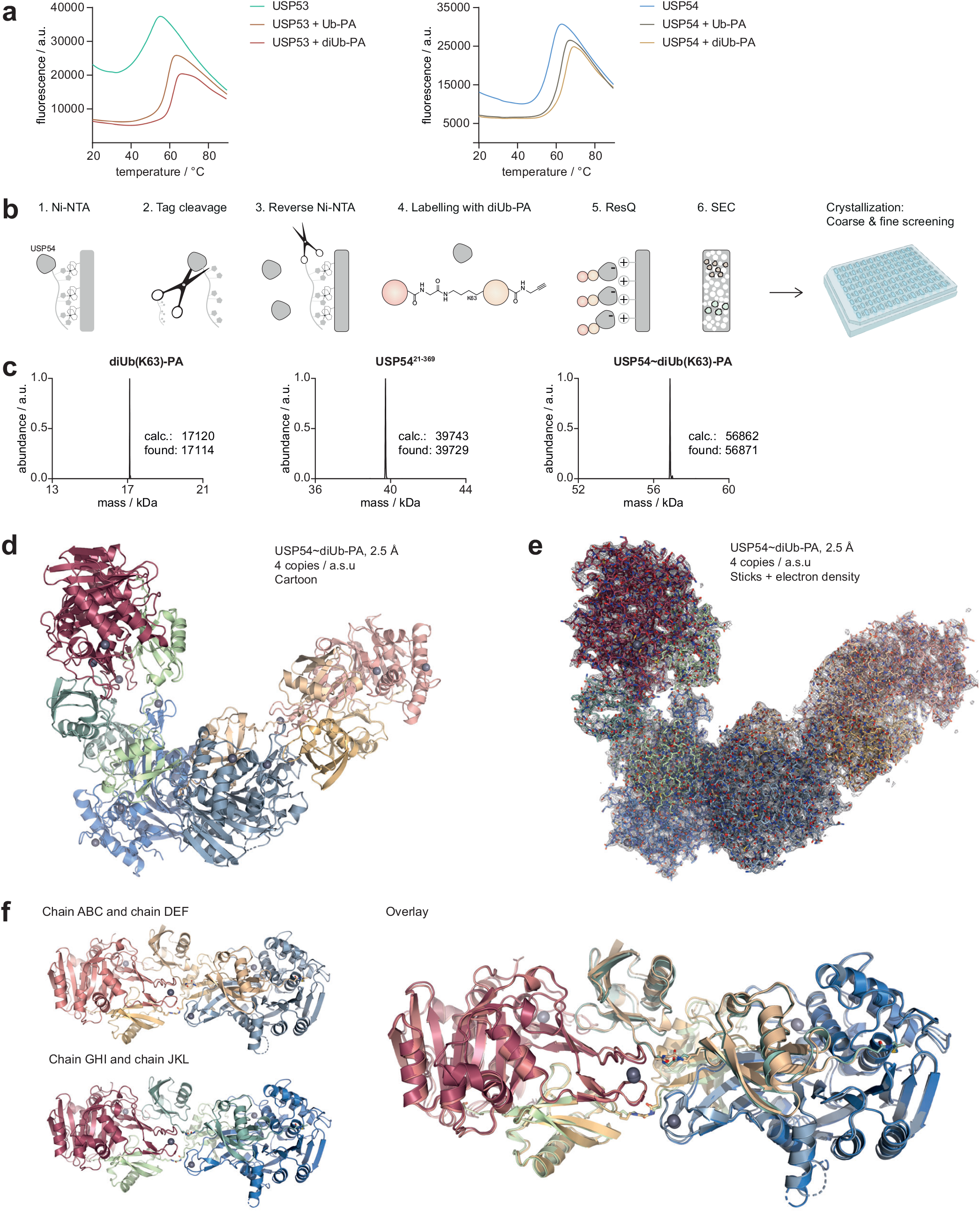
Assembly and crystallization of a USP54-diUb(K63)-PA complex. **a**. Fluorescence raw data of the protein stability assessment. Corresponding melting temperatures are shown in Fig. 4c. **b**. Assembly and purification strategy to obtain a pure USP54∼diUb(K63)-PA complex for crystallization. Of note, cleavage of the native isopeptide bond in the probe was suppressed by protein labeling at low temperature and with an excess of probe over enzyme. **c**. Deconvoluted intact protein mass spectra of the K63-linked diUb-PA probe, USP54, and the USP54∼diUb(K63)-PA complex. Calculated and observed molecular weights are given in Dalton. **d**. Asymmetric unit (a.s.u.) of the crystal structure of USP54∼diUb-PA. All four copies of USP54∼diUb-PA are shown in cartoon representation. **e**. Asymmetric unit (a.s.u.) of the crystal structure of USP54∼diUb-PA as in panel d, all four copies are shown in stick representation overlayed with 2F_O_-F_C_ electron density contoured at 1 σ. **f**. Chains ABCDEF and chains GHIJKL, each consisting of 2x USP54∼diUb-PA, are shown individually and as an overlay in cartoon representation.

**Supplementary Fig. 5.**
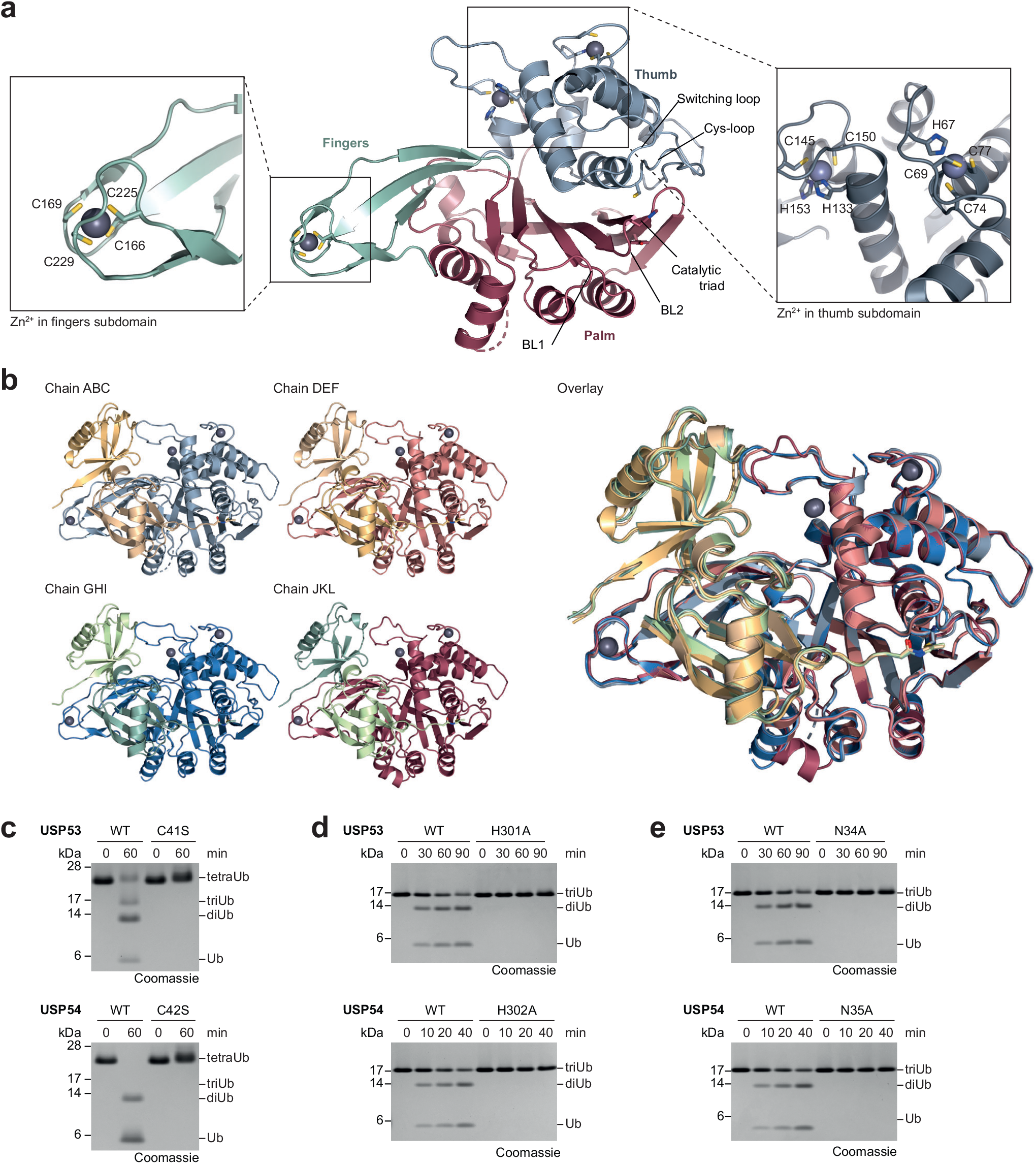
Analysis of the USP fold in the USP54∼diUb-PA structure. **a**. Typical fold of USP DUBs in the structure of USP54∼diUb-PA. The palm subdomain is colored red, the thumb subdomain blue and the fingers subdomain green. Additional structural elements including the blocking loops, the switching loop, the Cys-loop, the zinc ions, and the catalytic triad are annotated. Close-up views for the coordination of all zinc ions are shown. **b**. Chains ABC, DEF, GHI and JKL of the crystal structure of USP54∼diUb-PA, corresponding to the conformation in solution, are shown individually and as an overlay in cartoon representation. **c**. Gel-based cleavage assay assessing the catalytic cysteines. K63-linked tetraubiquitin chains were incubated with USP53^20-383^ (*upper*) or USP54^21-369^ (*lower*) and the respective catalytic cysteine mutants. Cleavage activity was analyzed by SDS-PAGE and Coomassie staining. **d**. Gel-based cleavage assay assessing the catalytic histidines. K63-linked triubiquitin chains were incubated with USP53^20-383^ (*upper*) or USP54^21-369^ (*lower*) and the respective catalytic histidine mutants for indicated time points. **e**. Gel-based cleavage assay as in d assessing potential oxy-anion hole residues.

**Supplementary Fig. 6.**
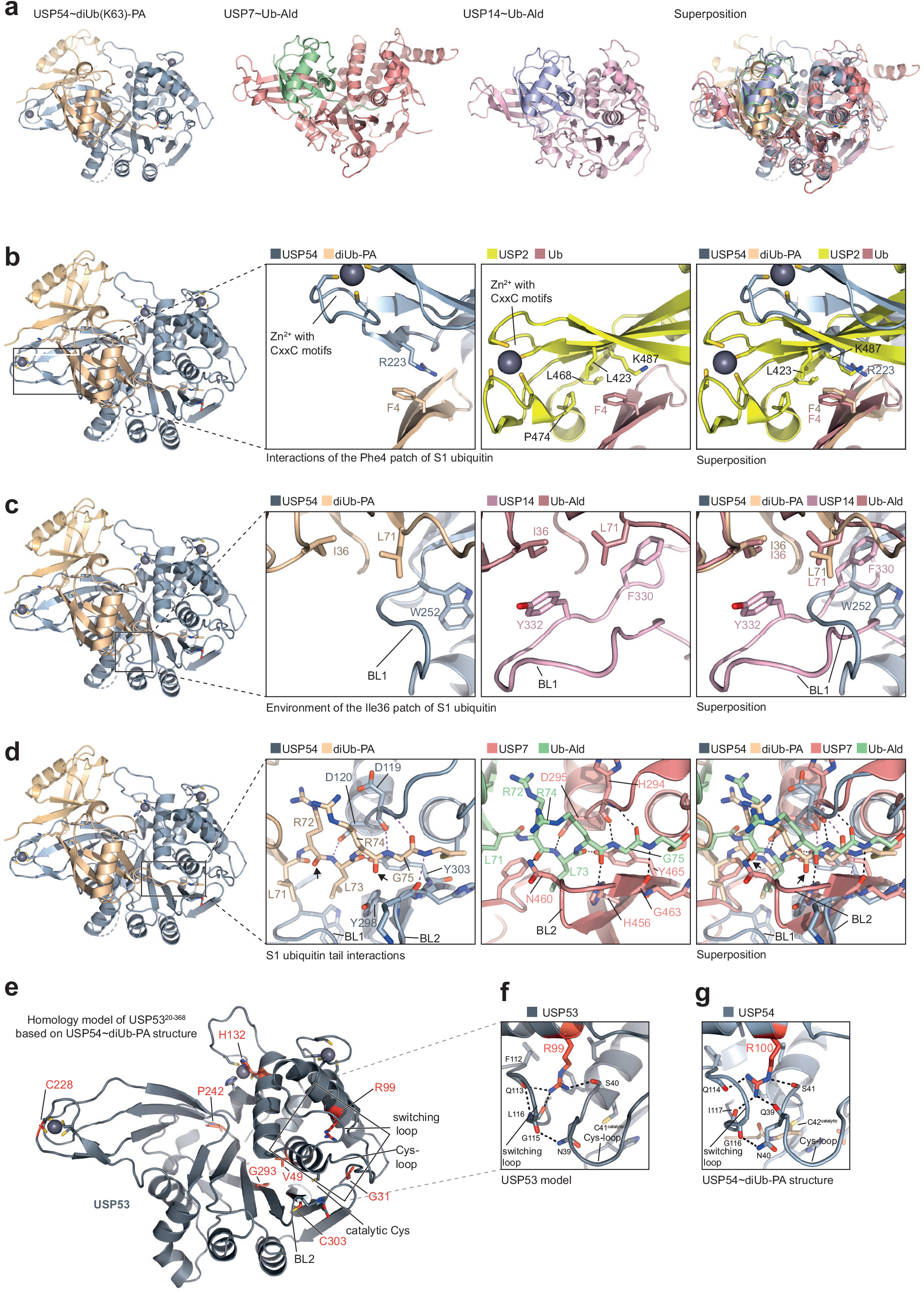
Analysis of the weakened S1 ubiquitin recognition in USP54. **a**. Cartoon representation of USP54∼diUb-PA as well as of representative USP DUB-ubiquitin aldehyde complexes (USP7, PDB: 1NBF; USP14, PDB: 2AYO). A superposition is shown, illustrating the distinct relative geometry of ubiquitin in the S1 site of USP54. **b**. Close-up view on the interaction of the Phe4 patch in the S1 ubiquitin with USP54 (*left*) and USP2 (PDB: 2IBI, *center*). A superposition (*right*) (aligned on ubiquitin) illustrates the different environment of Phe4 in USP54. **c**. Close-up view on the interactions of BL1 in USP54 (*left*) and in USP14 (PDB: 2AYO, *center)* with the S1 ubiquitin. A superposition (*right*) based on the S1 ubiquitin is shown. **d**. Close-up view on the S1 ubiquitin C-terminal tail interactions with USP54 (*left*) and USP7 (PDB: 1NBF, *center*). Hydrogen bonds are illustrated by dotted lines in pink for USP54 and black for USP7. Arrows highlight the missing hydrogen bonds in USP54 in comparison to USP7, also visible in the superposition (*right*). **e**. Homology model of the USP53 catalytic domain, based on the USP54 crystal structure. Residues of disease-associated single amino acid mutations are shown as red sticks (see Fig. 2a). **f**. Close-up view on Arg99 in USP53 highlights its bridging interactions to the switching loop and the Cys-loop. **g**. Close-up view on the corresponding residue R100 of USP54 in the crystal structure of USP54∼diUb(K63)-PA.

**Supplementary Fig. 7.**
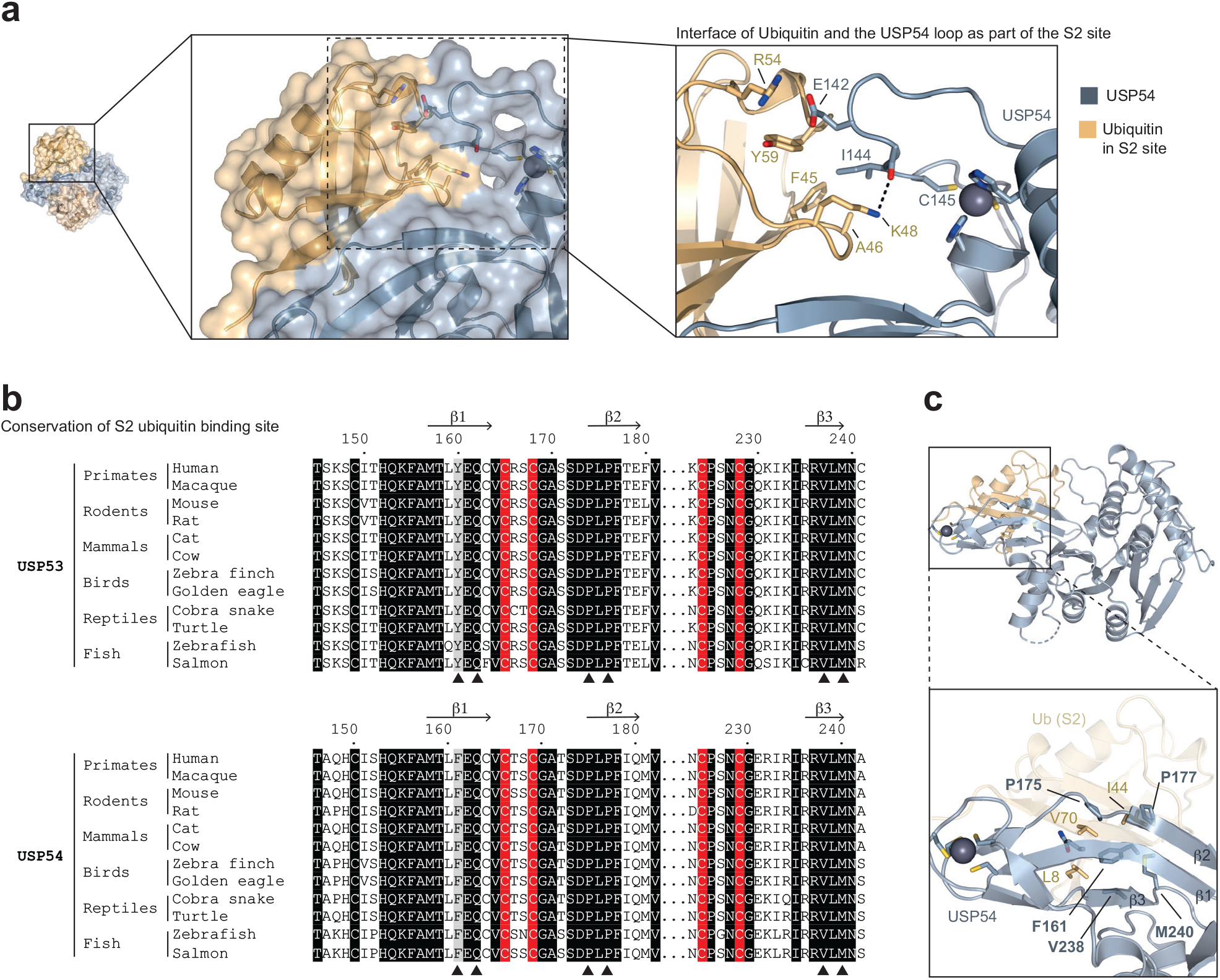
Conservation of S2 site on back of USP fingers in USP53 and USP54 orthologues. **a**. Close-up views on the S2 site. A loop in the thumb domain of USP54 above the S2 site forms hydrogen bonds to ubiquitin, flanked by a hydrophobic interaction between Ile144 of USP54 and Phe45 of ubiquitin. **b**. Sequence alignment of representative USP53 and USP54 orthologues showing the first three USP fingers beta sheets. Secondary structures are indicated according to the USP54∼diUb-PA structure, residue numbering is shown according to human sequences. Included species are human (*Homo sapiens*), Macaque monkey (*Macaca mulatta*), mouse (*Mus musculus*), rat (*Rattus norvegensis*), cat (*Felix catus*), cow (*Bos taurus*), zebra finch (*Taeniopygia guttata*), golden eagle (*Aquila chrysaetos chrysaetos*), Indian cobra (*Naja naja*), painted turtle (*Chrysemys picta bellii*), zebrafish (*Danio rerio*) and atlantic salmon (*Salmo salar*). Residues strictly conserved in all sequences are shown in black, the central aromatic residues are shown in grey. Cystein residues coordinating the zinc ion at the tip of the fingers are highlighted in red. Residues on the beta sheets with side chains pointing towards the backside / S2 ubiquitin binding site are highlighted with a black triangle. **c**. Coordination of the Ile44 patch in the S2 ubiquitin binding site. Cartoon representation of the USP54∼diUb-PA structure with close-up. Ubiquitin residues of the Ile44 patch are shown as sticks in yellow, residues highlighted with a black triangle in panel b are labeled as sticks in blue.

**Supplementary Fig. 8.**
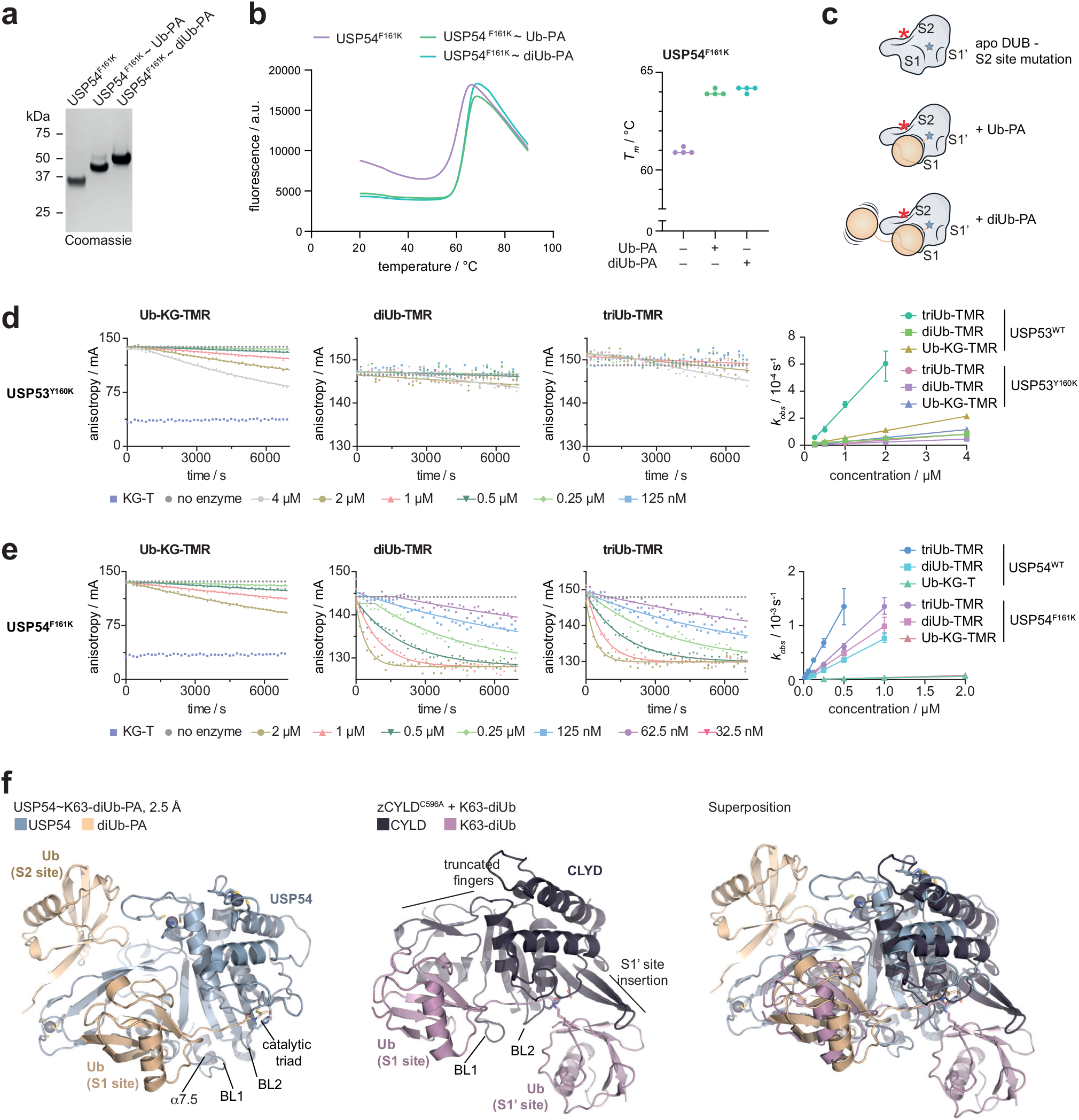
A cryptic S2 site mediates catalytic activity on longer ubiquitin chains. **a**. Purified samples of USP54 with mutated S2 site. USP54^21-369^ F161K, USP54^21-369^ F161K∼Ub-PA and USP54^21-369^ F161K∼diUb(K63)-PA were analyzed by SDS-PAGE and Coomassie staining. **b**. Protein stability assessment of S2 site-mutated USP54. Stability of samples from panel A was analyzed by thermal shift analysis. Fluorescence raw data are shown. Melting temperatures (*T*_*m*_) are plotted from technical replicates. **c**. Schematic of catalytic USP54 F161K DUB domains after reaction with Ub-PA or diUb-PA probes, illustrating ubiquitin engagement in samples used in panels A and B. S1’, S1 and S2 Ub binding sites are labelled. The mutated S2 site is depicted as a red star. **d**. Fluorescence polarization cleavage assays for USP53^20-383^ Y160K and indicated reagents. Fluorescence anisotropy was recorded over time and observed rate constants (*k*_*obs*_, shown as mean ± standard error) were plotted over enzyme concentrations. Values for WT protein were taken from Fig. 2e. **e**. Fluorescence polarization cleavage assays for USP54^21-369^ F161K and indicated reagents as described in panel d. **f**. Cartoon representation of USP54∼diUb-PA and CYLD in complex with K63-linked diUb (PDB: 3WXG). A superposition illustrates the different features enabling K63-linkage recognition in both enzymes by distinct mechanisms.

**Supplementary Fig. 9.**
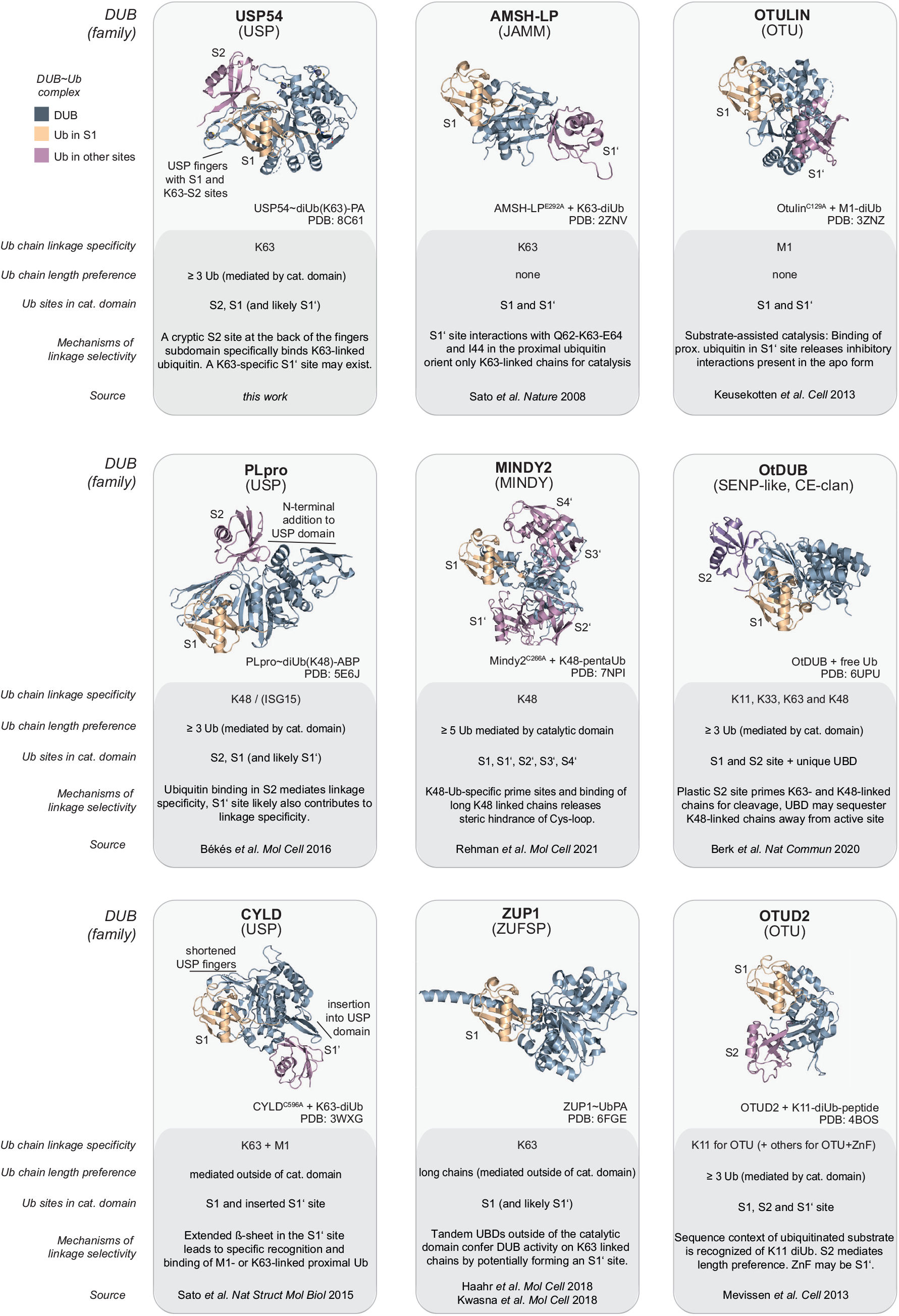
Mechanisms for polyubiquitin length- and linkage-specificity in DUBs.

## Notes

### Competing Interest Statement

The authors have declared no competing interest.

